# Stabilized gp120-specific CD4 for next-generation HIV-1 inhibitors

**DOI:** 10.64898/2026.03.24.713825

**Authors:** Adrian J. Bahn-Suh, Luis F. Caldera, Priyanthi N.P. Gnanapragasam, Jennifer R. Keeffe, Michael S. Seaman, Pamela J. Bjorkman, Stephen L. Mayo

## Abstract

HIV-1 Env’s gp120 subunit uses the T-cell coreceptor CD4 to enter host cells in a manner that prevents the evolution of host resistance by sharing the binding epitope with the footprint of CD4’s natural ligands, class II MHC proteins^1,2^. Consequently, CD4-containing biologics, such as CD4-Ig^3,4^ and derivatives^5–9^, benefit from this conserved relationship and are promising broad-acting anti-HIV-1 agents that are resistant to viral mutational escape^10^. However, these biologics suffer from short serum half-lives in humans^11,12^ and animals^3,13^, likely due to CD4’s poor thermostability^14^ and/or off-target class II MHC binding^15^. This latter property also warrants caution for CD4-containing biologics that could indiscriminately recruit Fc-dependent effector functions against uninfected cells and/or compete with host CD4 for class II MHC during T cell interactions with antigen-presenting cells. Here, we describe gp120-specific CD4 (gCD4), which exhibits enhanced thermostability and retains Env, but not class II MHC, binding. CD4-Ig variants incorporating gCD4 did not bind class II MHC on human B cells, displayed greater longevity in human tonsil organoid cultures, showed half-lives equivalent to therapeutic IgG antibodies in mice, and neutralized HIV-1 more broadly and potently compared to the original CD4-Ig molecules. Encouragingly, one variant neutralized 100% of a panel of clinically-relevant HIV-1 strains at titers correlating to infection prevention in humans, outperforming known broadly neutralizing antibodies^16,17^. Thus, gCD4 holds promise for the development of new CD4-containing biologics with best-in-class specificity, pharmacokinetic properties, and neutralization breadth and potency.

## Main

Broadly neutralizing antibodies (bNAbs) are clinically effective against HIV-1 but are susceptible to existing viral resistance or the development of viral mutational escape^16,18,19^. Addressing this shortfall with a pan-neutralizing and potent antibody that is resilient to HIV-1 escape has long been sought for therapeutic use. The pursuit of this goal led to the development of CD4-containing biologics such as CD4-Ig^3,4^, eCD4-Ig^7^, PRO-542^6^, and others^5,8,9,20^. These biologics fuse the two N-terminal domains of human CD4 (D1D2) to IgG Fc in various permutations and are attractive for therapeutic development because HIV-1 uses the gp120 subunit of its trimeric spike protein, Env, to bind to host cell CD4 and chemokine coreceptor to accomplish viral entry^21–25^. HIV-1 does this by first engaging CD4 through Env’s gp120 subunits, which then initiate conformational changes in Env to facilitate subsequent coreceptor binding and ultimately membrane fusion^26,27^. Therefore, a molecule such as eCD4-Ig, which contains two copies of CD4 and chemokine coreceptor mimetic peptides, is a broad neutralizer that is resilient to and constrains mutational escape—if HIV-1 were to evade neutralization by eCD4-Ig through mutations that reduce binding to CD4 or chemokine coreceptor, these mutations would also reduce HIV-1’s fitness. Evidence for this restrictive mechanism was demonstrated when viral swarms of HIV-1 under selective pressure by eCD4-Ig were unable to fully escape neutralization, and any viruses with partial resistance to eCD4-Ig exhibited markedly reduced infectivity^10^.

Various studies have shown CD4-containing biologics to prophylactically protect from HIV-1 infection and reduce viral loads during active infection *in vivo*, but these studies also showed evidence of poor pharmacokinetics^11,12,15^. For example, CD4-Ig and PRO-542 display half-lives of 2-4 days in mice^14^ and humans^11,12,15^, while typical IgGs have half-lives of 2-3 weeks in humans^28^. Many factors can affect serum half-life, but one property of CD4 that may be relevant to its pharmacokinetics is its low thermostability. Therapeutic monoclonal antibodies exhibit apparent melting temperatures (T_m_s) ranging from 60 - 90 °C^29^, but the CD4 domains used in CD4-containing biologics have much lower T_m_s (**Extended Data Table 1**). The poor thermostability of soluble CD4 constructs could lead to a greater propensity for the denaturation, aggregation, or misfolding of CD4 and contribute to short half-lives *in vivo*.

In addition, target-mediated drug disposition^15,30^ may play a role in the short serum half-lives of CD4-containing biologics. The natural ligands for CD4 are class II MHC proteins^31^ (MHC II), which are highly expressed on the surfaces of antigen-presenting cells (APCs) such as dendritic cells, macrophages, and B cells^32^. In a recent study relevant to the pharmacokinetics of MHC II-binding proteins, two T cell receptor (TCR)-like antibodies that either bound to all or to select peptide-MHC II complexes were investigated for their pharmacokinetic properties^33^. The antibody that bound to all peptide-MHC II exhibited rapid clearance, whereas the other exhibited more typical antibody pharmacokinetics. In a similar manner to the TCR-like antibodies, CD4-Ig may experience accelerated clearance through binding to MHC II.

We hypothesized that addressing these issues by stabilizing soluble CD4 and engineering it to preferentially bind HIV-1 gp120 and not MHC II would improve the pharmacokinetics of CD4-containing biologics. However, gp120 and MHC II share nearly identical binding surfaces on CD4—a feature that likely helps HIV-1 avoid the natural selection of protective CD4 polymorphisms^34^ as CD4 must maintain the ability to bind MHC II for a functional adaptive immune system. While beneficial for HIV-1, the shared binding interface on CD4 presented a considerable obstacle to our endeavor. Taking inspiration from the “divide and conquer” paradigm in software engineering (exemplified by the merge-sort algorithm^35^), we used computational and rational design methods to engineer CD4 variants that individually possess desired attributes before merging their properties in a final construct, which we refer to as gCD4 (for gp120-specific CD4). Compared to CD4-Ig molecules containing wild-type CD4, gCD4-Ig variants demonstrate better pharmacokinetics in tonsil organoid and humanized mice models and are more potent HIV-1 neutralizers, suggesting that the incorporation of gCD4 in future CD4-containing biologics would enhance their therapeutic properties.

### Electrostatic repulsions abrogated MHC II binding

We analyzed the co-crystal structures of the HIV-1 gp120- and human MHC II-CD4 complexes^1,2^ in PyMol^36^ to identify residues that could be substituted to selectively abrogate MHC II binding. The gp120-CD4 interface spans amino acid positions 23-64 in CD4, while the MHC II-CD4 interface spans amino acid positions 35-63 in CD4, a subset of the gp120 binding site (**Fig. 1a**). A critical difference between the two complexes is that gp120 is a monomer while MHC II is a heterodimer composed of MHC IIα and MHC IIβ. We aimed to exploit this difference by identifying CD4 residues unessential for binding gp120 and simultaneously crucial for binding MHC IIα or MHC IIβ. This analysis led to four observations. First, there are up to eight MHC IIβ interactions but only three MHC IIα interactions: CD4 residues R59, S60, and D63 interact with the corresponding MHC II α-chain residues T90, E88, and K176. Targeting the fewer number of MHC IIα interactions would be more feasible than targeting the MHC IIβ interactions. Second, MHC haplotypes are naturally diverse, but MHC IIα is invariant at these positions, circumventing the need to account for polymorphisms in this interface (**Fig. 1b**). Third, CD4 S60 and D63 are located at the periphery of the CD4-gp120 interface and have been shown in previous alanine-scanning experiments^37^ to be dispensable for gp120 binding. Therefore, mutating these residues should have minimal effects on CD4’s affinity for gp120 and should preserve the ability of CD4-containing biologics to restrict HIV-1 mutational escape. Finally, these positions are near the N- and C-termini of a short α-helix in CD4. An α-helix possesses a macrodipole that is more positively charged near its N-terminus and more negatively charged near its C-terminus. Thus, offsetting this macrodipole by introducing a negatively charged residue at S60 and a positively charged residue at D63 would likely have a favorable effect on CD4 thermostability^38^, as well as introduce unfavorable electrostatic interactions with the MHC IIα chain E88 and K176 (**Fig. 1b**). Accordingly, we set out to investigate the effects of alanine, aspartate, glutamate, or lysine substitutions at CD4 positions 60 and 63, hypothesizing that the charged substitutions would have the strongest effects in abolishing MHC II binding.

**Fig. 1:**
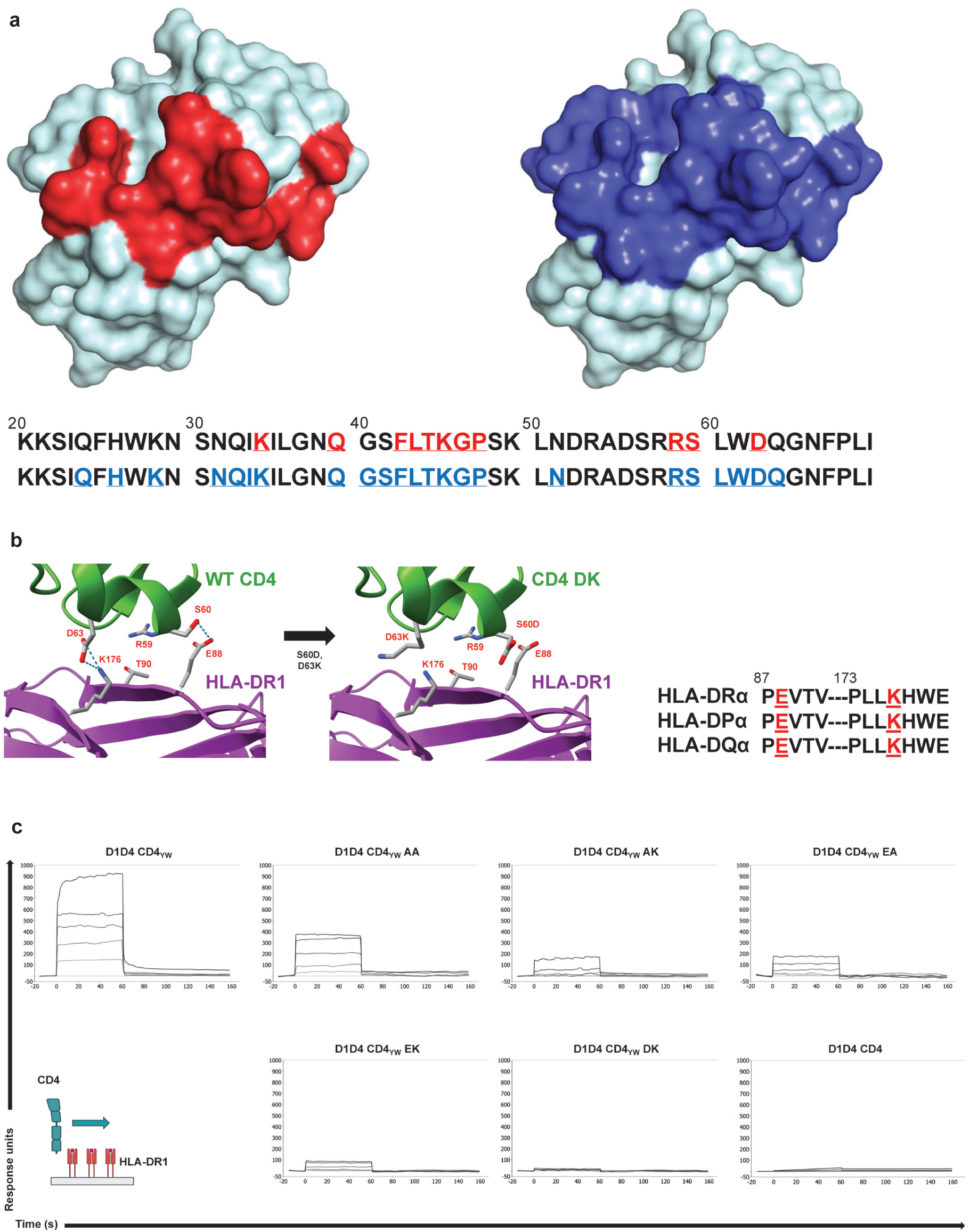
Design of MHC II binding knockdown substitutions for CD4. **a,** Surface representation of the D1 domain of CD4 (residues 1-100) in pale cyan with the MHC II-interacting residues in red on the left and the gp120-interacting residues in blue on the right (PDB: 1GC1). The CD4 sequence is shown in text starting from position 20 and ending at position 70. The MHC II-interacting residues are underlined in red, and below them the gp120-interacting residues are underlined in blue. **b,** Cartoon representation of the interface of CD4 (top; green) and HLA-DRα (bottom; purple) and stick representation of side chains in close proximity (PDB: 3S4S). Dashed lines indicate S60 and D63 of wild-type CD4 (left) interacting with E88 and K176 of HLA-DRα. Modeling the S60D and D63K substitutions in CD4 DK (middle) show unfavorable electrostatic interactions with HLA-DRα. The sequences of HLA-DRA, HLA-DPA, and HLA-DQA are aligned and shown on the right (aa 87-91 and 173-179), and E88 and K176 are underlined in red. **c,** SPR sensorgrams of HLA-DR1 and variants of the full ectodomain of CD4. Biotinylated HLA-DR1 was immobilized on the streptavidin-conjugated surface and multi-cycle kinetics was performed using CD4 at various concentrations (100 μM to 1.23 μM in 3-fold dilutions). Injections were performed starting with SPR buffer and proceeded from the lowest concentration to the highest concentration. No regeneration was performed between injections. A cartoon of the SPR conditions is shown in the bottom left. CD4 variants with Q40Y and T45W are denoted in subscript, and S60 and D63 substitutions are indicated by the final paired letters (e.g. “CD4_YW_ AA” contains Q40Y, T45W, S60A, and D63A).

Normally, CD4 acts together with a TCR to recognize peptide-MHC II complexes^39^, but CD4 alone demonstrates low^40^ (>2 mM K_D_) monomeric affinity for MHC II *in vitro*. Hence, we used a previously reported affinity-matured CD4^2^, CD4_YW_, to evaluate the effects of our designed CD4 substitutions using surface plasmon resonance (SPR)^41^. The Q40Y/T45W substitutions in CD4_YW_ only improve interactions with MHC IIβ and would not be expected to confound the results of the MHC IIα-binding knockdown substitutions. Since CD4 S60R and D63R increased binding to MHC II in the prior CD4 affinity-maturation^2^, arginine was not included as a potential amino acid substitution at CD4 position 63. With these considerations, we performed surface plasmon resonance (SPR) binding assays using the full ectodomain CD4_YW_ D1-D4 containing various S60/D63 substitutions against MHC II (**Fig. 1c**). As hypothesized, the charged substitutions displayed greater reductions in MHC II binding than the alanine substitutions, in rank order of strongest to weakest binding: CD4_YW_ AA > CD4_YW_ AK ∼ CD4_YW_ EA > CD4_YW_ EK > CD4_YW_ DK. Remarkably, CD4_YW_ DK’s reduced affinity for MHC II appeared to be equivalent to that of the negative control—CD4 that lacks the high-affinity Q40Y/T45W substitutions and therefore has negligible binding to MHC II in SPR assays.

### Surface and core optimizations improve stability

In parallel, we explored optimizing the surface exposed and buried core amino acids of CD4 to engineer improved thermostability. Analysis of the surface properties of D1D2 was performed in PyMol^36^ and the TRIAD physics-based protein design platform to identify design targets for mutagenesis. Our structure-guided analysis identified a hydrophobic patch in D2 that consisted of the side chains L109, L151, V175, and L177 (**Fig. 2a)**. These residues would normally be partially buried due to contacts with CD4 D3 in the full CD4 ectodomain^42^; however, they become fully exposed when CD4 is truncated to L177 as in CD4-Ig. We then performed *in silico* mutational energy score calculations^43,44^ in TRIAD to determine amino acid substitutions that would stabilize this exposed hydrophobic patch (**Extended Data Fig. 1)**. Nine of the most favorable predictions were expressed and screened for thermostability using differential scanning fluorimetry^45^ (DSF) to determine their T_m_s. The best performing set of substitutions, L109K, L151K, V175I, and L177D (KKID), improved the T_m_ of D1D2 by 6 °C (**Fig. 2b**). We investigated the mechanism for this improvement by modeling L109K, L151K, and L177D, observing that the substituted residues could form a network of salt bridges at the exposed C-terminus of CD4 D1D2 (**Fig. 2a**). We also tested a known space-filling A55V substitution^5^ in D1 that improved the T_m_ of D1D2 by 8 °C (**Fig. 2b**). When the A55V and KKID substitutions were combined, the resulting VKKID variant stabilized D1D2 by 14 °C (**Fig. 2b**). It is likely that the two sets of substitutions additively improved the T_m_ by independent mechanisms: the A55V substitution stabilized the hydrophobic core of the D1 domain, and the KKID substitutions stabilized the hydrophilic surface of the D2 domain. DSF measurement results for all variants are compiled in **Extended Data Table 1**.

**Fig. 2:**
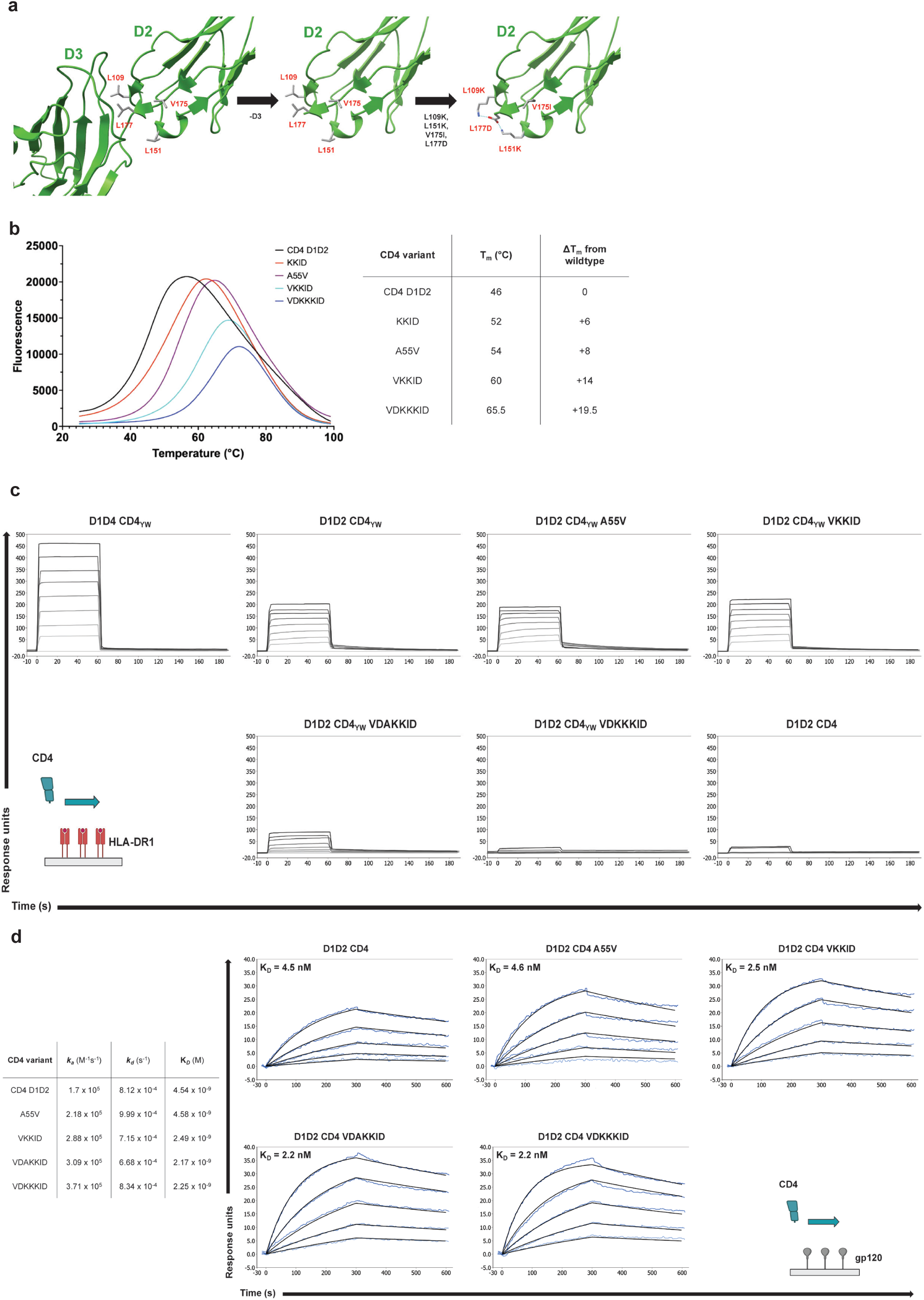
MHC II knockdown and thermostabilizing substitutions preserve CD4’s affinity for gp120. **a,** Cartoon representation of D2 and D3 (left; green) from the full ectodomain of CD4 (PDB: 1WIP) and the side chains of L109, L151, V175, and L177 as sticks. Truncating the D3 domain fully exposes the previously partially buried hydrophobic residues (middle). Modeling the L109K, L151K, V175I, and L177D substitutions (right) shows a newly formed stabilizing network of electrostatic interactions in the KKID variant. **b,** DSF data of CD4 variants (left) and their melting temperatures in tabular form (right). **c, d** SPR sensorgrams of CD4 variants against HLA-DR1 (**c**) and gp120 (**d**). **c,** Biotyinlated HLA-DR1 was immobilized on streptavidin-conjugated surfaces and CD4 variants were injected for multi-cycle kinetics as in **Fig. 1c** at various concentrations (100 μM to 0.78 μM in 2-fold dilutions). **d**, Full-length BG505 gp120 was immobilized using primary amine coupling and CD4 variants were injected for multi-cycle kinetics at various concentrations (31.25 nM to 1.95 nM in 2-fold dilutions). The gp120 surfaces were regenerated with 10 mM glycine pH 3.0 between injections. Experimental data (blue) were globally fitted using a 1:1 Langmuir binding model (black) and the kinetic rate constants are reported in tabular form (left). Cartoons of the SPR conditions are shown for **c** (bottom left) and **d** (bottom right).

### gCD4 is thermostable and gp120-specific

After identifying the S60 and D63 substitutions for reducing MHC II binding, we merged these with the VKKID substitutions to make thermostable D1D2 constructs with reduced affinity for MHC II. DSF of the resulting VDKKKID variant indicated that the S60D and D63K substitutions improved thermostability as hypothesized (**Fig. 2b**). We then performed SPR with our D1D2 constructs against gp120 and MHC II to confirm that binding to gp120 was retained and only binding to MHC II was knocked down. SPR with MHC II showed that only CD4_YW_ D1D2 variants with substitutions at S60 and D63 had reduced affinity for MHC II, and the results reinforced that S60D and D63K were the most disruptive substitutions (**Fig. 2c**). Furthermore, the SPR with gp120 confirmed that all evaluated CD4 D1D2 variants exhibited single-digit nM affinity for gp120, consistent with previously reported values^46,47^ (**Fig. 2d**). From these data, we concluded that our targeted substitutions did not disrupt CD4’s ability to bind gp120.

Additional mutational screening of KKID revealed that the V175I substitution was unnecessary and that L177E was slightly more thermostable than L177D, perhaps due to glutamate’s longer side chain facilitating more favorable salt-bridge formation with the substituted residues in L109K and L151K. We thus defined gp120-specific CD4, or gCD4, as the set of D1D2 substitutions containing A55V, S60D, D63K, L109K, L151K, and L177E (VDKKKE). Compared to wild-type D1D2, gCD4 exhibited minimal evidence of thermal denaturation at physiologic temperatures, an improved T_m_ (66.5 °C) by >20 °C, and yielded nearly 50-fold more protein from mammalian cell expressions (**Extended Data Table 1 and Extended Data Fig. 2**).

### gCD4-Ig inherited the properties of gCD4

Having developed gCD4 with increased thermostability and diminished binding to MHC II, we asked whether these qualities would be inherited if gCD4 were introduced into the CD4-Ig and eCD4-Ig Fc-fusion molecules. DSF of the CD4-Ig and eCD4-Ig variants indicated that gCD4-Ig (T_m_=64.5 °C) and e-gCD4-Ig (T_m_=62.5 °C) were 10 °C and 9 °C more thermostable than CD4-Ig and eCD4-Ig, respectively (**Extended Data Fig. 3**). Surprisingly, gCD4-Ig and e-gCD4-Ig had lower melting temperatures than their soluble gCD4 counterpart, likely because the Fc domains undergo thermal denaturation before the gCD4 domains: human Fc initiates its first melting transition around 60-65 °C^48^, consistent with our observations. We next repeated SPR with MHC II using CD4-Ig variants to investigate how avidity would influence the binding interactions. Similar to the behavior of monomeric CD4, introducing the S60D and D63K substitutions to CD4_YW_-Ig in the form of gCD4_YW_-Ig greatly reduced binding to MHC II (**Extended Data Fig. 4**). After establishing that gCD4-Ig showed diminished MHC II binding by SPR, we investigated how our CD4-Ig variants would behave in more biologically relevant contexts.

We characterized CD4-Ig’s ability to engage MHC II *ex vivo* through flow cytometry of primary human tonsillar leukocytes^49^ to account for the surfaces of APCs being densely coated with MHC II^50^. We visualized CD4-Ig by introducing an Avitag^51^ to the C-terminus of the Fc, enabling biotinylation and detection with fluorophore-conjugated streptavidin (**Fig. 3a**). Clinical trials of PRO-542, a CD4-IgG2 Fc-fusion molecule, achieved serum concentrations of ≥1 μM with a single 10 mg/kg dose and reported rapid clearance of PRO-542^11,12^. Mimicking those conditions, tonsil cells were labeled using 1 μM (100 μg/mL) of the CD4-Ig variants (**Figs. 3b-g**). Flow cytometry with CD4-Ig variants containing the affinity-enhancing Q40Y/T45W substitutions indicated that CD4_YW_-Ig labeled the majority of CD19+ tonsillar B lymphocytes (**Fig. 3d**). Comparatively, gCD4_YW_-Ig labeled a smaller proportion of tonsillar B lymphocytes (**Fig. 3e**). Without the affinity-enhancing substitutions, CD4-Ig also labeled a small proportion of B lymphocytes (**Fig. 3f**), an unexpected result due to wild-type CD4’s weak monomeric affinity for MHC II^40^. Notably, gCD4-Ig weakly labeled tonsillar B lymphocytes (**Fig. 3g**), and the measured fluorescence was only slightly above background levels seen with the negative control condition (nonbiotinylated Avitagged CD4_YW_-Ig) (**Fig. 3c**). We observed an identical staining pattern using tonsil cells from an unrelated donor (**Extended Data Fig. 5**), supporting the generalizability of the S60D and D63K substitutions in abolishing MHC II binding irrespective of MHC haplotype.

**Fig. 3:**
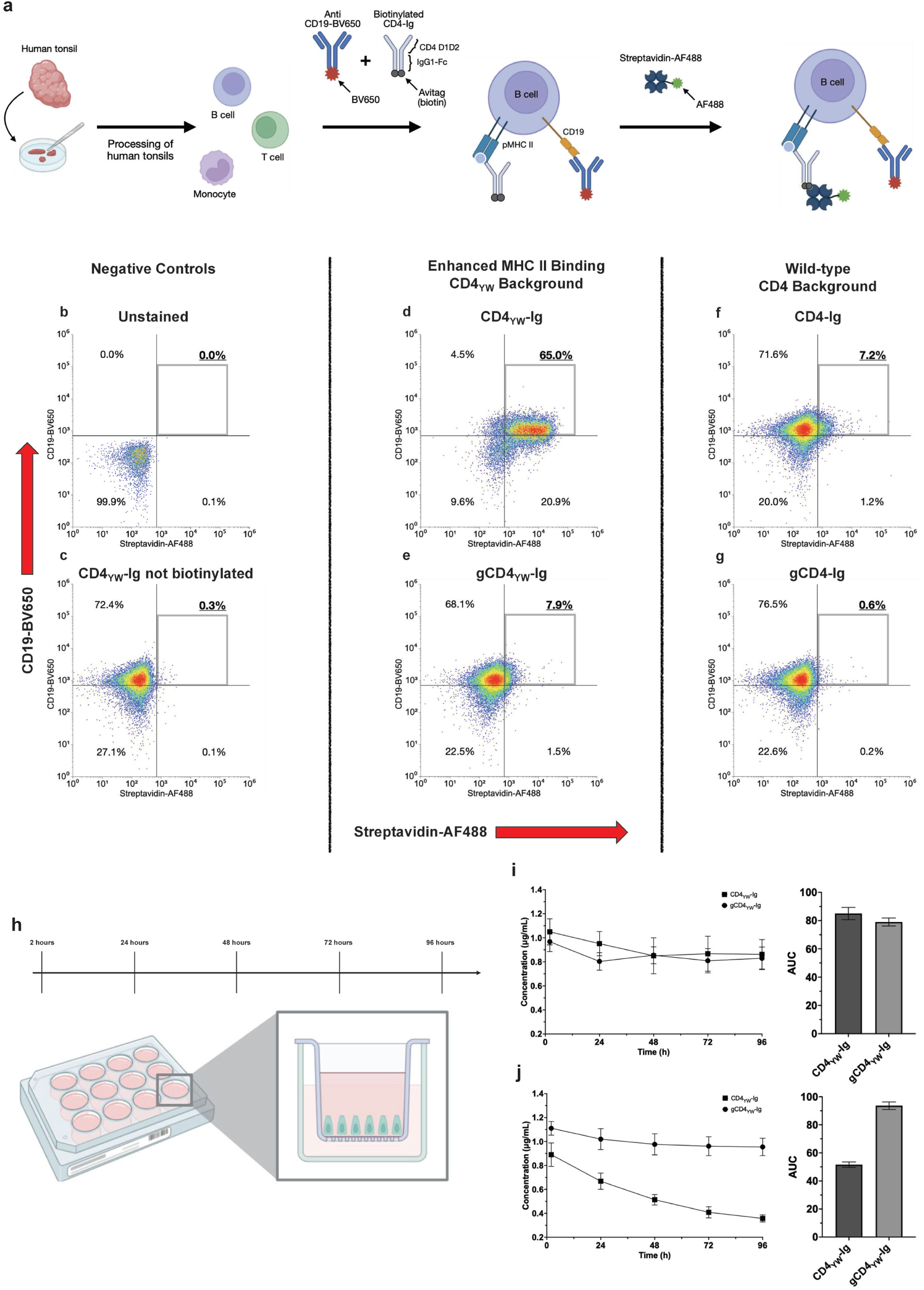
gCD4-Ig persists in tonsil organoid cultures due to reduced MHC II binding. **a,** Diagram of the flow cytometry experimental set-up. Mononuclear leukocytes were isolated by Lymphoprep gradient density separation, aliquoted, and stored frozen. Aliquots were thawed as needed and samples were prepared with live-dead staining and Human TruStain FcX Fc receptor blocking solution. The cells were then labeled with anti-CD19 conjugated to Brilliant Violet 650 and biotinylated CD4-Ig variants. After removing excess antibody, the cells were labeled with streptavidin conjugated to Alexa Fluor 488 and analyzed by flow cytometry. **b-g,** Flow cytometry plots of tonsillar cells. CD19 signal is plotted on the y-axis and streptavidin signal is plotted on the x-axis. The plots were gated on viable singlet cells, and the fluorescence thresholds were determined using the negative controls (unstained cells and stained cells labeled with unbiotinylated Avitagged CD4_YW_-Ig). The proportion of double-positive cells for CD19 and CD4-Ig are indicated in the top right quadrant. **h**, Graphic of the tonsil organoid culture experimental set-up. Five to six million tonsil cells were aliquoted in 12-well transwell plates with each well containing a 1 mL outer reservoir of media. 20 μL from the outer reservoirs were sampled at 2 hours, 24 hours, 48 hours, 72 hours, and 96 hours post-transwell seeding. The outer reservoirs were mixed by pipetting prior to each collection and the timepoints were stored at -80 °C. **i, j,** Time-course measurements (left) of the concentration of CD4_YW_-Ig and gCD4_YW_-Ig incubated in tonsil culture media without tonsil cell seeding (**i**) or with tonsil cell seeding (**j**) and their area under the curves (right). Concentrations were determined by ELISA in technical triplicates using the respective proteins as standards. Each point is an average of three independent transwell samples and the error bars are s.d. Variants containing the Q40Y and T45W substitutions are denoted in subscript. CD4_YW_-Ig for **i** and **j** additionally contain the thermostabilizing A55V, L109K, L151K, and L177E substitutions. gCD4 constructs for **e, g, i**, and **j** contain A55V, S60D, D63K, L109K, L151K, and L177E substitutions. Donor #664 was used for the data in **b-g** and donor #669 was used for the data in **i** and **j**.

After observing that CD4-Ig measurably binds to cell surface MHC II, we tested our hypothesis that target-mediated drug disposition plays a role in CD4-Ig pharmacokinetics by employing a human tonsil organoid model^49^ (**Figs. 3h-j**). We estimated that a leukocyte:CD4-Ig mass ratio of 1000:1 would simulate a typical 15 mg/kg CD4-Ig infusion dose for a 70 kg reference adult male^52^, which translates to a concentration of 10 nM (1 μg/mL) in the tonsil organoids (**Extended Data Fig. 6**). To compensate for the 100-fold lower concentration compared to our flow cytometry experiments, we used CD4_YW_-Ig constructs at 10 nM (**Extended Data Figs. 7d and 7e)** as a first approximation for the behavior of CD4-Ig at 1 μM **(Extended Data Figs. 7b and 7c**). Tonsil organoids from a single donor were cultured for four days in the presence of B-cell activating factor (BAFF), which promotes B lymphocyte survival and surface MHC II expression^49,53^. When CD4_YW_-Ig proteins with and without the MHC II-abrogating DK substitutions were incubated in only culture media, there was no difference in protein loss over time (**Fig. 3i**). However, in the presence of tonsil cells, the construct with intact MHC II binding underwent accelerated clearance from the culture, whereas the construct with abrogated MHC II binding did not **(Fig. 3j**). These data suggest that MHC II binding contributes to accelerated CD4-Ig turnover in the tonsil organoids.

After characterizing gCD4-Ig’s behavior with MHC II-presenting tonsil cells, we evaluated neutralization of a global representative 12-strain panel of HIV-1 pseudoviruses^54^ by gCD4-Ig. Although gCD4 did not demonstrate improved binding affinity to gp120 by SPR, its inclusion in CD4-Ig and eCD4-Ig improved the half-maximal inhibitory concentrations (IC_50_s) of the precursor molecules: gCD4-Ig (geometric mean IC_50_ 0.74 μg/mL) and e-gCD4-Ig (geometric mean IC_50_ 0.035 μg/mL) were 5.6 and 7.7-fold more potent than CD4-Ig and eCD4-Ig, respectively (**Fig. 4a**). CD4-Ig and gCD4-Ig were both not polyreactive in a baculovirus polyreactivity assay^55^ (**Extended Data Fig. 8**), suggesting that non-specific interactions did not account for the improved neutralization. Instead, we hypothesize that the potency increases may be due to improvements in the conformational stability of gCD4 rather than higher affinity for gp120.

**Fig. 4:**
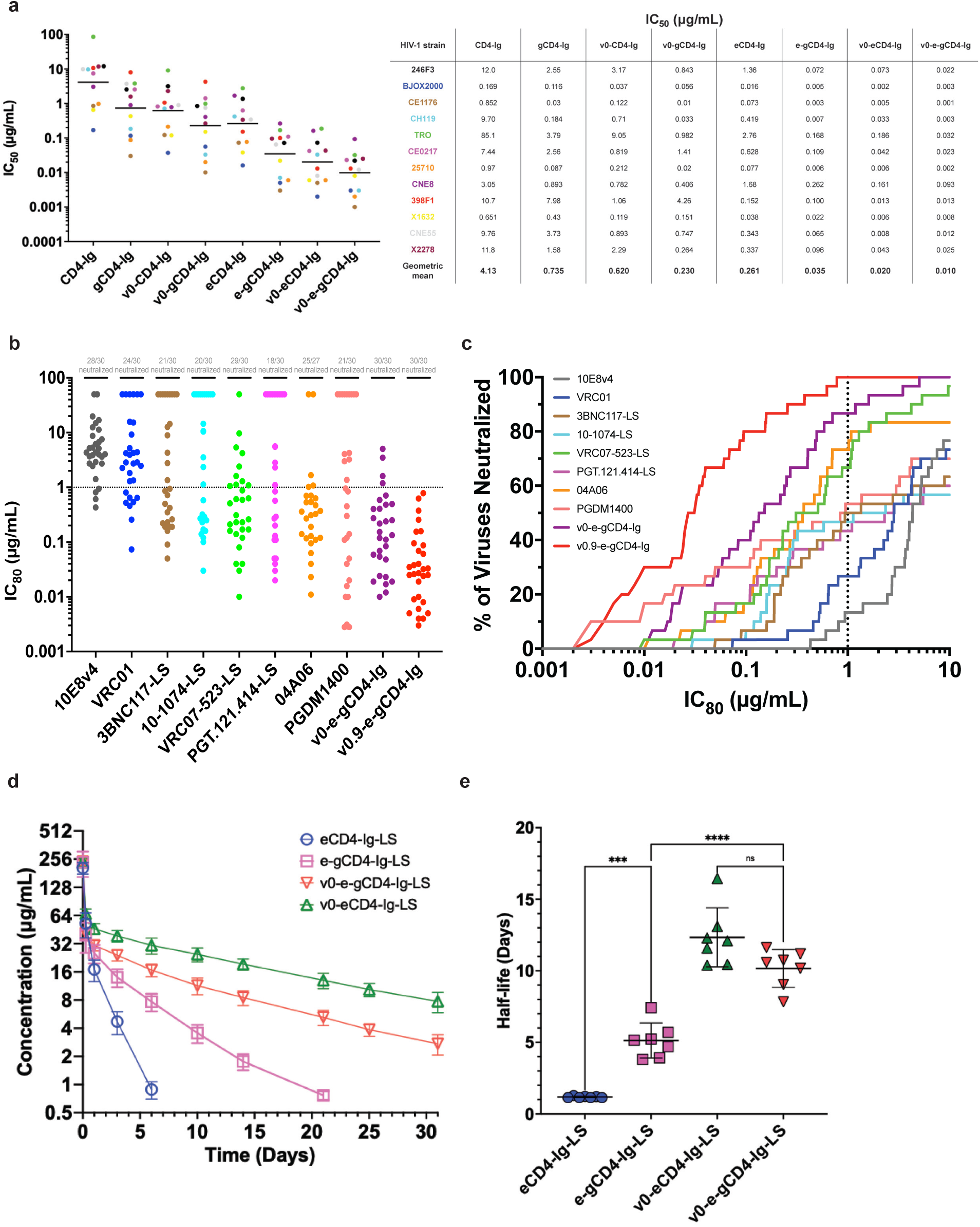
CD4-Ig incorporating gCD4 display enhanced HIV-1 neutralization and have longer half-lives in humanized mice. **a,** IC_50_ data of CD4-Ig variants against the global 12-strain panel in Tzm-bl cell assays as a scatter plot (left) and table (right). Each point represents the average of technical duplicates for a given virus strain and the black bars indicate the geometric means. **b,** IC_80_ data of HIV-1 bNAbs and e-gCD4-Ig variants against a panel of 30 viruses from the AMP trials. Non-neutralized viruses are shown as points at 50 μg/mL. **c,** Coverage curves of the viruses from the AMP trials using the data in **b**. Each curve indicates the fraction of the virus panel that is 80% neutralized at a given antibody concentration. The protective IC_80_ threshold from the AMP trials is signified by the dotted lines in **b** and **c** at 1 μg/mL. Neutralization data of the bNAbs were obtained from the CATNAP database. **d**, PK data of eCD4-Ig-LS variants in SCID hFcRn transgenic mice. Each group contained seven mice and the averages of the plasma concentrations are plotted. **e,** Scatter plot of the terminal half-lives of eCD4-Ig-LS variants. Half-lives were determined from individual mice and the average half-lives are presented as black bars. Statistical significance was determined by one-way ANOVA using Tukey’s multiple comparisons (***: p < 0.001; ****: p < 0.0001; ns: not significant). Error bars in **d** and **e** are s.d.

### gCD4-Ig variants have therapeutic potential

During these latest experiments, a v0-CD4-Ig containing the CD4 substitutions G6A, S23N, A55V, V128L, R134G, N164D, K167T, and V168L was demonstrated to be more thermostable than wild-type CD4-Ig^14^. Since gCD4 only has one substitution in common—A55V—we tested whether the two sets of substitutions were compatible with each other. We first investigated the determinants of v0-CD4-Ig’s thermostability by expressing its CD4 D1D2 domains and performing DSF. v0-CD4 displayed a T_m_ of 78 °C, with most of its thermostability contributions coming from the hydrophobic core substitutions G6A, A55V, V128L, and V167L (v0_core_, T_m_ of 72 °C). The combination of the v0-CD4 and gCD4 substitutions, v0-gCD4, had the highest T_m_ of 80 °C, and we again observed that Fc-fusion constructs had melting temperatures that were lower than the most stable CD4 domains (**Extended Data Table 1**), supporting our hypothesis that the Fc’s thermal denaturation occurs before the stabilized CD4 domains. We next introduced the Q40Y/T45W substitutions into v0-CD4 and tested its binding to MHC II by SPR. Similar to the previous SPR results, v0-CD4_YW_ bound to MHC II, and the introduction of the S60D and D63K substitutions in v0-CD4_YW_ DK and v0-gCD4_YW_ significantly reduced MHC II binding (**Extended Data Fig. 9**). We obtained equivalent results with v0.9-CD4_YW_ and v0.9-gCD4_YW_, which contained additional R59K/K90R substitutions that enhance affinity for gp120^14^ **(Extended Data Fig. 9**). Taken together, these data suggest that S60D and D63K are necessary and sufficient to reduce binding to MHC II, independent of the extensive amino acid substitutions in v0-CD4.

Since v0-gCD4 combined the favorable properties of v0-CD4 and gCD4, we evaluated whether v0-gCD4 would similarly improve CD4-Ig’s neutralization potency. CD4-Ig and eCD4-Ig molecules containing v0-gCD4 demonstrated the highest potency against the 12-strain panel compared to those containing gCD4 or v0-CD4 (**Fig. 4a**). The Antibody Mediated Prevention (AMP) trials^16^ had determined that protective neutralization titers correlate to *in vitro* IC_80_ values less than 1 μg/mL; thus, we asked if our most potent variants could neutralize below this threshold by evaluating v0-e-gCD4-Ig and v0.9-e-gCD4-Ig against a panel of 30 AMP trial pseudoviruses. Compared to known highly potent bNAbs, all of which failed to provide complete coverage against the panel, both v0-e-gCD4-Ig and v0.9-e-gCD4-Ig potently neutralized all viruses. The additional gp120 affinity-enhancing R59K and K90R substitutions in v0.9-e-gCD4-Ig improved potency over v0-e-gCD4-Ig as previously reported^14^, and v0.9-e-gCD4-Ig neutralized all viruses below the 1 μg/mL threshold (**Figs. 4b** and **4c** and **Extended Data Table 2**). From these results, v0.9-e-gCD4-Ig demonstrated superior breadth and potency against AMP viruses than current therapeutic bNAbs, suggesting that passive immunization with v0.9-e-gCD4-Ig could provide protection against circulating HIV-1 strains.

Finally, we asked how the enhanced thermostability of v0-gCD4 affected the pharmacokinetics of eCD4-Ig by performing pharmacokinetic studies of eCD4-Ig variants containing the half-life enhancing LS mutations^56^ using SCID human FcRn transgenic mice^57,58^. These mice are deficient in MHC II due to impaired lymphocyte development^58^ and low levels of APC activation^59^, enabling us to minimize MHC II binding as a confounding variable in our half-life estimations. Our studies revealed that eCD4-Ig-LS had a short half-life of ∼1 day despite containing the LS mutations, but e-gCD4-Ig-LS and v0-e-gCD4-Ig-LS demonstrated improved terminal half-lives of ∼5 and ∼10 days (**Figs. 4c and Fig. 4d**), respectively. v0-eCD4-Ig had the longest terminal half-life of ∼12 days, but this difference was not statistically significant compared to v0-e-gCD4-Ig (p=0.19), and both molecules achieved half-lives in mice on par with known therapeutic bNAbs^60^. The net charge of an antibody is known to affect its pharmacokinetics^61^, although electrostatics may play a minor role in our results as e-gCD4-Ig has a longer half-life than eCD4-Ig despite having more positively charged surface residues. We instead speculate that the relative rank order of the eCD4-Ig variants’ half-lives in the SCID mice may be most reflective of their respective CD4 domains’ thermostabilities. From these findings, we conclude that thermostabilizing the CD4 domains improved the pharmacokinetics of eCD4-Ig in SCID human FcRn transgenic mice.

## Discussion

Here, we provide evidence that CD4’s affinity for MHC II can be abolished while retaining CD4’s affinity for HIV-1, paving the way for the clinical development of CD4-containing biologics incorporating gCD4. Using CD4-Ig as a proof-of-principle, our data demonstrated that we have produced HIV-1 entry inhibitors with pharmacokinetic properties and neutralizing capabilities that surpass those of therapeutic bNAbs. Encouragingly, our most potent variant neutralized all tested pseudoviruses from the AMP trials at titers indicative of protective efficacy, a feat that would normally require a mixture of different bNAbs^17^. Other CD4-containing biologics, such as bNAb-CD4 fusions^8^, CD4 nanoparticles^20^, and multimeric CD4-Fc constructs^5^ are likely to benefit from the inclusion of gCD4 in their future constructions. Finally, our computational and rational design approaches can be reasonably applied to other proteins to engineer their biophysical and biochemical properties.

Our observation that CD4-Ig binds to cellular MHC II raises several considerations for the clinical use of CD4-containing biologics as it remains unknown how such molecules might interact with human physiology. A previous study indicated that soluble monomeric CD4 did not acutely inhibit CD4+ T cell activation *in vitro*^62^, but it is possible that multimeric molecules such as CD4-Ig would compete for MHC II engagement and interfere with CD4+ T cell activation and/or development due to greater avidity. It is also not known if CD4-Ig would indiscriminately mediate Fc receptor-dependent cytotoxicity against MHC II-containing cells. These unknowns motivated precaution in the clinical formulation of PRO-542, which contained four copies of CD4 fused to IgG2 Fc to weaken Fc effector functions^6^. However, these concerns are now mitigated by gCD4’s near-complete specificity for HIV-1, which should enable safer implementation of CD4-Fc fusions with intact effector functions. Fc effector functions are key features of protective antibodies, allowing them to kill virally infected cells through mechanisms such as antibody-dependent cellular cytotoxicity (ADCC). For example, it was shown that eCD4-Ig with IgG1, but not IgG2, Fc can mediate ADCC of HIV-1-infected cells *in vitro*^63^. Fc effector functions are also implicated in the ability of bNAbs to prolong control of viremia via the vaccinal effect^64^. Due to these advantages, biologics with IgG1 Fc would be preferred for developing curative therapeutic regimens for people living with HIV-1. In addition to treating active infection, it has been shown that eCD4-Ig delivered by AAV can act like an HIV-1 vaccine and provide years-long protection from HIV-1 infection in infant rhesus macaques^65^. In the context of AAV gene therapy, reducing MHC II binding may become a more salient consideration for potentially lifelong eCD4-Ig expression.

Limitations of our study include the functional relevance of our tonsil and mouse experiments in the context of human physiology and the translatability of our *in vitro* neutralization results to *in vivo* efficacy. First, our flow cytometry experiments likely underestimate how much CD4-Ig binds to MHC II *in vivo*. The rapid dissociation kinetics of CD4 and MHC II as seen in our SPR data imply that a large proportion of CD4-Ig would readily disengage from the tonsillar B cells during washing and that the binding at equilibrium would be higher than what we have recorded. Second, the effect size of MHC II-mediated clearance remains unknown physiologically, although our tonsil organoid data suggest that MHC II binding results in measurably faster clearance. Transgenic mice expressing fully human MHC II may be an appropriate model to test this variable. Third, the preparation of tonsil organoids results in a limited composition of mononuclear white blood cells that are predominantly B and T lymphocytes. Macrophages express MHC II and make up a much larger proportion of the human immune system by mass than any other individual white blood cell^52^. Therefore, macrophages may significantly contribute to the clearance of CD4-Ig *in vivo* through the reticuloendothelial system. Since spleens contain large numbers of macrophages, switching to spleen-derived organoids may address this limitation^49,52^. Fourth, we did not test whether CD4-Ig binding to MHC II has functional relevance, such as investigating B and T lymphocyte activation in the tonsil organoids longitudinally. Finally, although our e-gCD4-Ig variants displayed excellent neutralization breadth and potencies, we did not evaluate if they prevent or treat HIV-1 infection *in vivo*. However, the comparatively less potent eCD4-Ig was shown to prevent HIV-1 infection in mice and rhesus macaques^13,65,66^; thus, it is likely that our molecules will behave similarly although further studies are required to investigate the efficacies of e-gCD4-Ig variants. While more challenges remain, gCD4 demonstrates great potential to develop superior CD4-containing biologics to provide safe, durable, and broad protection and/or treatment against HIV-1.

## Materials and Methods

### DNA constructs and vectors

Human codon-optimized CD4 sequences with a GS linker and C-terminal 6xHis tag were designed and ordered as gBlocks from IDT using IDT’s codon optimization tool. The gBlocks were cloned into the vectors pTT5 (National Research Council of Canada) or p3BNC through Gibson assembly^67^ using NEBuilder HiFi DNA Assembly Master Mix (New England Biolabs) according to manufacturer recommendations. DNAs were transformed into chemically competent Mix and Go! DH5α *E. coli* (Zymo Research) and grown on LB agar plates containing 100 μg/mL carbenicillin. Plasmids were isolated from *E. coli* using Zippy Plasmid Miniprep Kits (Zymo Research) and verified through Oxford nanopore sequencing (Plasmidsaurus).

CD4-Ig and eCD4-Ig constructs were designed as previously reported. gBlocks were ordered from IDT and assembled in p3BNC as above. All CD4-Ig constructs contained identical mouse VH leader sequences. The sequence for human TPST2 was obtained from Uniprot (Accession ID: O60704) and cloned into p3BNC with its native Golgi signal sequence. For Avitagged proteins, the 15 amino acid Avitag sequence GLNDIFEAQKIEWHE was added to the C-terminus of the Fc. The sequence for *E. coli* biotin ligase was assembled in p3BNC as above and contained a mouse VH leader sequence and a C-terminal KDEL endoplasmic reticulum retention motif. Plasmid DNAs for mammalian cell transfection were prepared from *E. coli* using Macharey-Nagel midiprep and maxiprep kits.

The pTT5 plasmid containing His-tagged full-length BG505 gp120 was obtained from the Bjorkman lab.

### Protein expression and purification

Gibco Expi293F cells were transfected with 1 μg of DNA/mL of cells using the Gibco ExpiFectamine transfection kit according to manufacturer recommendations. Four days post-transfection, cultures were centrifuged and supernatants were filtered through 0.45-μm filters. For His-tagged proteins, Pierce disposable columns were packed with HisPur Ni-NTA resin (Thermo Scientific), rinsed with water, and equilibrated with Gibco 1X Dulbecco’s PBS with no magnesium and no calcium, pH 7.4 (DPBS). Clarified supernatant was applied to equilibrated Ni-NTA resin under gravity flow. Columns were washed with ten column volumes of wash buffer (DPBS, 15 mM imidazole) and eluted with two to three column volumes of elution buffer (DPBS, 300 mM imidazole). Eluted proteins were concentrated using Amicon spin concentrators and further purified via FPLC using a Superdex 200 Increase 10/300 GL column equilibrated with DPBS at 4°C. Fractions containing the desired protein were pooled, concentrated, filtered with a 0.22 μm filter, and stored at 4°C. Protein concentrations were determined by measuring absorbance at 280 nm on a NanoDrop (Thermo Fisher Scientific). Extinction coefficients were calculated by inputting the amino acid sequences of the mature proteins in Expasy ProtParam.

Expression and purification of CD4-Ig and eCD4-Ig was performed as previously described^14^. Plasmids containing human TPST2 and eCD4-Ig were co-transfected at a 1:4 DNA mass ratio of TPST2:eCD4-Ig. Expression of biotinylated CD4-Ig was performed in an identical manner using a 1:4 ratio of BirA:Avitagged CD4-Ig. Purification of Fc-containing proteins was performed using HiTrap MabSelect SuRe columns (Cytiva) connected to a peristaltic pump according to manufacturer recommendations. Columns were washed with ten column volumes of 1X DPBS and eluted using IgG elution buffer (Pierce) into 1 M Tris-HCl buffer, pH 9 (NC9904344; Boston Bioproducts). Eluted proteins were further purified via FPLC as above.

### Computational design of CD4

For *in silico* predictions of thermostabilizing substitutions, the coordinates of CD4 D1D2 were extracted from cocrystal structures of CD4 D1D2 bound to gp120 using PyMOL (PDBs: 1GC1, 2NY1, 2NY2)^1,68^. Water molecules were not included. Rosetta^43^ and PHOENIX^44^ force fields were used in TRIAD to separately standardize and analyze the extracted CD4 structures. Monte Carlo optimization with sampling was performed without covalent terms on fixed backbones from initial rotamers using the expanded Dunbrack rotamer library^69^ and floating nearby residues. Sequence logos of the predicted sequences were generated using WebLogo (https://weblogo.berkeley.edu/). A subset of *in silico* predictions were selected for experimental testing based on the frequency of each residue at each position as well as the relative rank order of the unique combinations.

### Differential scanning fluorimetry

DSF assays were performed by preparing triplicate samples in DPBS to a final concentration of 5 to 10 μM of protein and 20X Sypro Orange dye (Invitrogen). A CFX96 Touch Real-Time PCR Detection System (Bio-Rad) was used to measure the fluorescence of the samples by heating from 25.0°C to 99.0°C in 0.5°C increments and 30 second equilibration times. Background subtraction was performed using the mean fluorescence of triplicate samples containing only buffer and dye. The data were exported and plotted in GraphPad Prism 11. Apparent melting temperatures were defined as the inflection point of the first derivative of the fluorescence data with respect to time.

### Surface Plasmon Resonance

Analyte proteins for SPR injection were prepared as described above with one modification: the FPLC buffer was substituted for 1X HBS-EP+ (150 mM NaCl, 10m HEPES pH 7.4, 2mM EDTA, 0.05% Tween-20) prepared from 20X HBS-EP+ buffer (Teknova). Multi-cycle kinetics were performed at 25°C in a Bruker Sierra SPR Pro-32 instrument. SPR running buffer was 1X HBS-EP+ and SPR wash buffer consisted of 0.05% Tween-20. The ligand for SPR experiments in **Figs. 1c** and **2c** was biotinylated monomeric HLA-DR1 loaded with influenza A hemagglutinin peptide 306-318 (PKYVKQNTLKLAT), obtained from the NIH Tetramer Core Facility. The ligand for subsequent SPR experiments was biotinylated monomeric HLA-DR1 loaded with SARS-CoV-2 S peptide 873-888 (GAALQIPFAMQMAYRFF), also obtained from the NIH Tetramer Core Facility. Biotinylated ligands were diluted to 100 nM in running buffer and directly immobilized on a Bruker Biotin-Tag Capture Sensor to achieve response units (RU) of approximately 500-1,000. No additional protein was immobilized on reference flow cells as they already contained streptavidin. Three start-up injections of buffer preceded analyte injections. Analyte samples were prepared in 2-fold or 3-fold dilution series in running buffer and injected at a flow rate of 30 uL/min followed by a disassociation phase. CD4 and CD4-Ig exhibited extremely rapid dissociation kinetics and required no surface regeneration between cycles. Analyte injections were performed in the order from lowest concentration first to highest concentration last. For SPR of CD4 and gp120, full-length BG505 gp120 was diluted to 100 nM in 10 mM sodium acetate pH 4.5 and immobilized on a Bruker high-capacity amine chip to approximately 400-900 RU according to manufacturer recommendations. M8a-3 IgG, an anti-SARS-CoV-2 antibody, was equivalently coupled to reference flow cells^70^. Injections were performed with similar conditions as above and included a regeneration step with 10 mM glycine pH 3.0. SPR sensorgrams for all experiments were double subtracted from the reference and buffer injections. Experimental data for SPRs with gp120 were globally fit to a 1:1 Langmuir model using the Bruker Sierra Analysis software.

### Baculovirus assay

Polyreactivity was assessed using an ELISA-based method to assay non-specific binding to baculovirus particles as described^55,71^. Briefly, a 1% solution of baculovirus particles in 100 mM sodium bicarbonate buffer, pH 9.6 was adsorbed to a 384-well Maxisorp ELISA plate overnight at 4°C. After blocking with 0.5% bovine serum albumin (BSA) in PBS, CD4-Ig samples and controls were added to the blocked assay plate and incubated for 3 hours at room temperature. The plate was washed with PBS and an HRP-conjugated anti-IgG secondary antibody (Southern Biotech 2015-05) was added and incubated for one hour at room temperature. Plates were washed with PBS and bound IgG was detected by measuring luminescence at 425 nm after addition of SuperSignal ELISA Femto Maximum Sensitivity Substrate (ThermoFisher).

### Pseudovirus Neutralization

HIV-1 pseudovirus neutralization assays were performed and analyzed as described^72,73^, either in-house^74^ or through the Collaboration for AIDS Vaccine Discovery (CAVD) core neutralization facility. Briefly, HIV-1 pseudoviruses were generated by cotransfecting HEK 293T cells with an HIV-1 Env expression plasmid and a replication-defective backbone plasmid^72,73^. Supernatants were harvested at 72 hours, filtered through a 0.45 um sterile filter, then titered using TZM-bl target cells, which express HIV-1 receptors CD4 and CXCR5^72^. CD4-based inhibitors were evaluated in duplicate with a 5-fold, 8-point dilution series of the inhibitor. The CD4-based inhibitor was incubated with pseudovirus for one hour at 37°C then added to TZM-bl target cells. After 48 hours at 37°C, cells were lysed with BriteLite luminescence substrate and luminescence was measured to evaluate the amount of infection at each concentration of inhibitor. The percent neutralization was calculated and plotted as a function of concentration. IC_50_ and IC_80_ values, the concentrations at which half-maximal or 80% neutralization were observed, were derived using a nonlinear regression analysis using the Antibody Database^75^. These values were exported and plotted in GraphPad Prism 11.

### Tonsil Organoids

De-identified human tonsils were obtained from the Cooperative Human Tissue Network and processed as previously reported^49^. Briefly, tonsils were sectioned and manually homogenized into suspension. Mononuclear leukocytes were isolated by Lymphoprep density gradient separation, washed with complete media (RPMI with glutamax, 10% Fetal Bovine Serum [FBS], 1X nonessential amino acids, 1X sodium pyruvate, 1X penicillin–streptomycin, 1X Normocin [InvivoGen] and 1X insulin/selenium/transferrin cocktail [Gibco]), and frozen in 1 mL aliquots of 50-100 million cells in FBS + 10% dimethyl sulfoxide (DMSO). Frozen aliquots were stored at -140 °C. Tonsil cells were prepared for culturing by thawing in a 37 °C water bath and resuspending in 10 mL of prewarmed thaw media. Resuspended cells were centrifuged at 500 x g for 10 minutes at 4 °C and the thaw media was aspirated. Pelleted cells were resuspended in prewarmed complete media, and the number of viable cells was determined with Trypan blue using a hemocytometer. The cell suspension was then adjusted to a density of ∼60 million cells/mL using complete media. Culture plates were prepared by adding 1 mL of prewarmed complete media supplemented with 1 μg/mL of human B cell-actvating factor (BAFF; BioLegend) and, if applicable, CD4-Ig proteins to 12-well plates (Corning). 12 mm diameter Millicell culture plate inserts (MilliporeSigma) were placed into each well and 100 μl of cell suspension was plated into the inserts. For media-only wells, 100 μl of complete media was added to the inserts. Cultures were incubated at 37 °C and 5% CO_2_ with humidity. Aliquots of culture media were taken from the exterior reservoir after 2 hours, 24 hours, 48 hours, 72 hours, and 96 hours and frozen at -80 °C for later analysis. The remaining volumes of blank wells filled with complete media were tracked using a micropipette to estimate evaporation over time in the cultures and cross-verified with the experimental wells at the end of the experiments. ELISA signals were multiplied with a correction factor for evaporation at each time point, accounting for approximately 50 μL of evaporation every 24 hours. Timepoints and the area under the curves were plotted using GraphPad Prism 11.

### Flow Cytometry

Frozen tonsil aliquots were thawed in a 37 °C water bath and centrifuged for 3 minutes at 1,000 x g at 4 °C. The media was discarded, and the cells were washed by resuspending the cell pellet with DPBS. The cells were pelleted and stained with Zombie Violet Fixable Viability Kit (423113; BioLegend) according to manufacturer recommendations in DPBS. After viability staining, cells were washed once with FACS buffer (DPBS, 0.1% Bovine Serum Albumin [BSA; 50-178-1669; Sigma Aldrich]) and cellular Fc receptor was blocked with Human TruStain FcX (422302; BioLegend) according to manufacturer recommendations in FACS buffer. Approximately 1-2 million cells were aliquoted into individual samples of 100 μL of FACS buffer and incubated with 3 μL of Brilliant Violet 650 anti-human CD-19 antibody conjugate (302238; BioLegend) and 1 μM purified Avitagged CD4-Ig proteins for 1 hour on ice protected from light. Samples were pelleted and resuspended in 50 μL of FACS buffer containing streptavidin-Alexa Fluor 488 conjugate (S32354; Invitrogen) diluted 1:50 (final concentration 40 μg/mL). After 30 minutes of incubating on ice protected from light, the samples were washed once with 200 μL of FACS buffer and kept pelleted on ice until the time of measurement. Samples were resuspended in 500 μL of FACS buffer, filtered through 70 μm Flowmi pipette tip strainers (H13680-0070; SP Bel-Art), and immediately analyzed on a Sony SH800S Cell Sorter. Compensation matrices were calculated using unstained and single-stain controls and 50,000 to 100,000 total events were recorded for each sample. Cells were gated for singlets, viability, and the detection of Brilliant Violet 650 and streptavidin-Alexa Fluor 488. Flow cytometry plots were generated using Floreada.

### ELISA

Determination of the CD4-Ig concentration in the tonsil organoid cultures was performed by capture ELISA. Clear, flat-bottom, high bind 96-well polystyrene plates (Corning) were coated overnight at 4 °C with 50 μL of mouse anti-human CD4 antibody, clone RPA-T4 (300502; BioLegend), diluted to 2 μg/ml in 0.1 M NaHCO_3_, pH 9.6. Wells were emptied by dumping into a sink and then blotted on clean paper towels. Coated wells were blocked with 300 μL of blocking buffer (DPBS, 3% BSA, 0.1% Tween-20) at room temperature (RT) for 2 hours. Frozen aliquots of the tonsil culture media samples were thawed and diluted 1:100 into blocking buffer. The blocked plates were then dumped and blotted, followed by application of 50 μL of the diluted tonsil culture media samples in triplicate. Standard curves of purified CD4-Ig variants were applied on the same plate in triplicate using 4-fold serial dilutions in blocking buffer starting from a top concentration of 1 μg/mL. The plates were incubated at room temperature (RT) for 2 hours and washed three times with washing buffer (DPBS, 0.1% Tween-20). HRP-conjugated goat anti-human IgG Heavy Chain Superclonal antibody (A56862; Invitrogen) was prepared by diluting 1:10,000 in blocking buffer, and 50 μL of secondary antibody was applied to each well and incubated at RT for 30 minutes. After three washes, 50 μL of 1-Step Ultra TMB-ELISA substrate (34028; Thermo Scientific) was applied to each well and incubated at RT for ten minutes protected from light. The reaction was stopped by adding 50 μL of 1 N HCl to each well and absorbance at 450 nm was measured with a plate reader. The concentration of CD4-Ig in the tonsil samples was estimated by interpolating from standard curves using a four-parameter nonlinear regression model in GraphPad Prism 11. Correction factors from the measured evaporation were applied to the estimated concentrations.

### Murine pharmacokinetics

All experimental work relating to mice was performed by The Jackson Laboratory. Four groups of SCID hFcRn mice (JAX strain# 018441) at 6-8 weeks of age were intravenously given 10 mg/kg of eCD4-Ig-LS variants. Whole blood was collected at 5 minutes, 6 hours, 1 day, 3 days, 6 days, 10 days, 14 days, 21 days, 25 days, and 31 days from the tail vein and processed to plasma before being stored at -80 °C. eCD4-Ig-LS concentrations were assayed by capture ELISA using an anti-CD4 capture antibody and a HRP-conjugated goat anti-human secondary antibody as above. PK analysis was performed using Phoenix WinNonLin 8.5.2.4.

## Acknowledgments

We thank Lisa Wagar and Sam Kim (UC Irvine) for help in establishing tonsil organoid systems, Jost Vielmetter and the Caltech Protein Expression Center for assistance with protein expression and purification, Jost Vielmetter and Harry Gristick for discussions about SPR, Rebecca Hill (Kaiser Permanente School of Medicine) for help with performing power analyses in designing mouse experiments, the NIH Tetramer Core Facility for providing biotinylated HLA-DR1 monomers, and Mike McCune (Gates Foundation) and members of the Mayo and Bjorkman labs for helpful discussions. This work was supported by the Bill and Melinda Gates Foundation INV-057196 (P.J.B.) and INV-036842 (M.S.S.).

## Author Contributions

A.J.B., L.F.C., P.J.B., and S.L.M. conceived the study and P.J.B. and S.L.M. provided funding support. A.J.B., L.F.C., P.J.B., and S.L.M. designed experiments. M.S.S. and P.N.P.G. performed HIV-1 neutralization assays, J.R.K. performed polyreactivity assays. A.J.B. and L.F.C. performed and analyzed all other experiments. A.J.B., L.F.C., P.J.B., and S.L.M. wrote the manuscript.

## Extended Data

**Extended Data Table 1:**
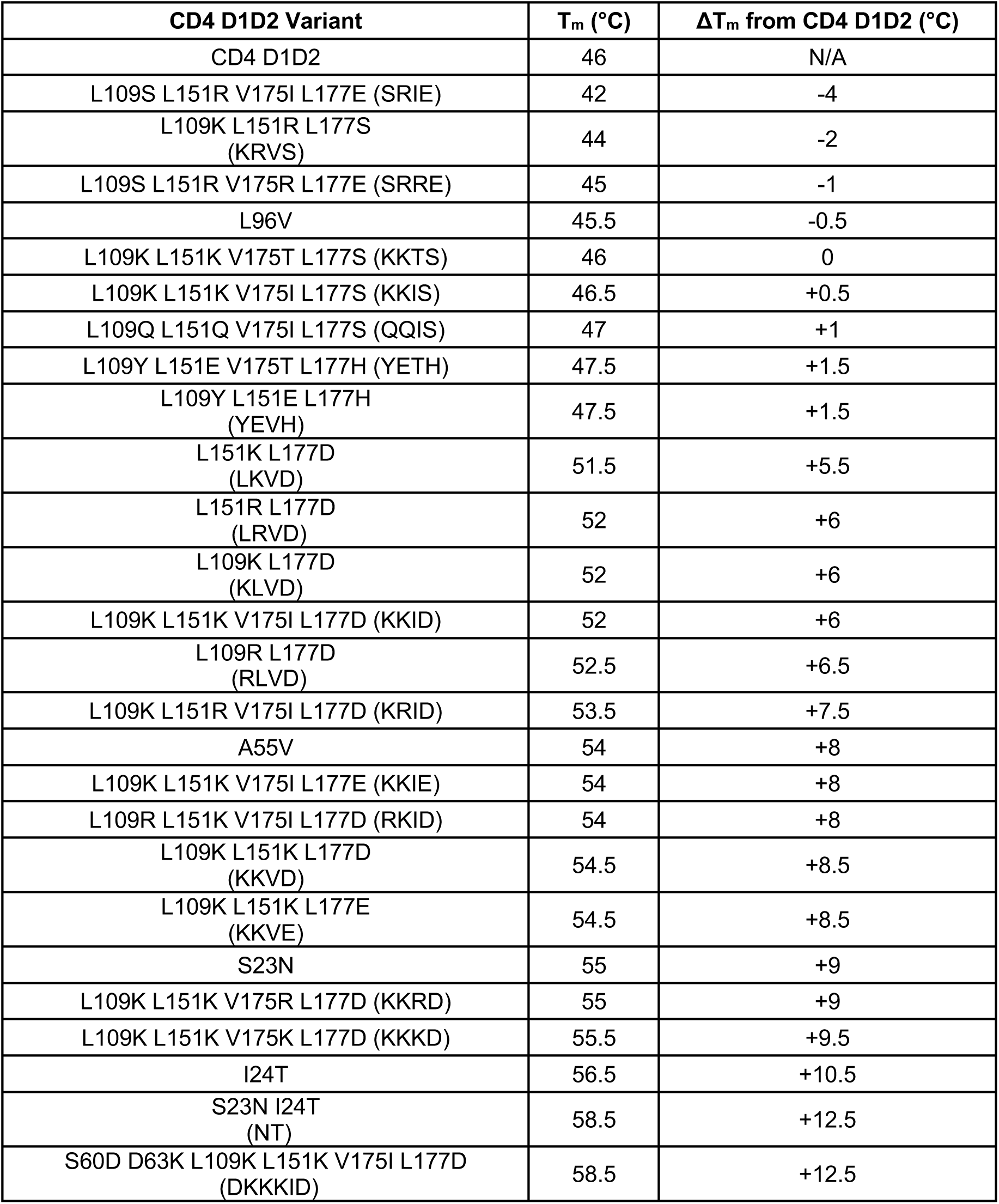
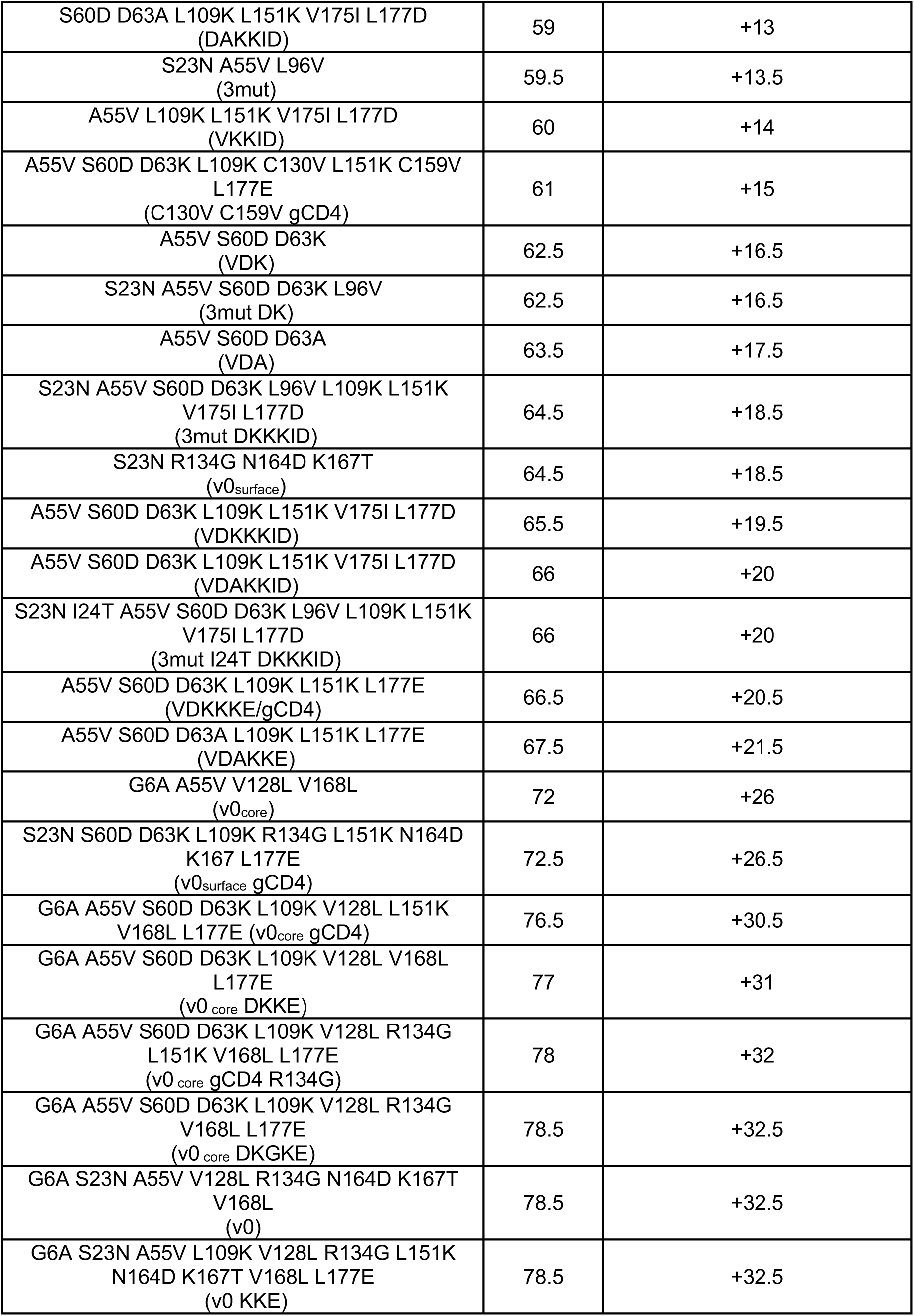
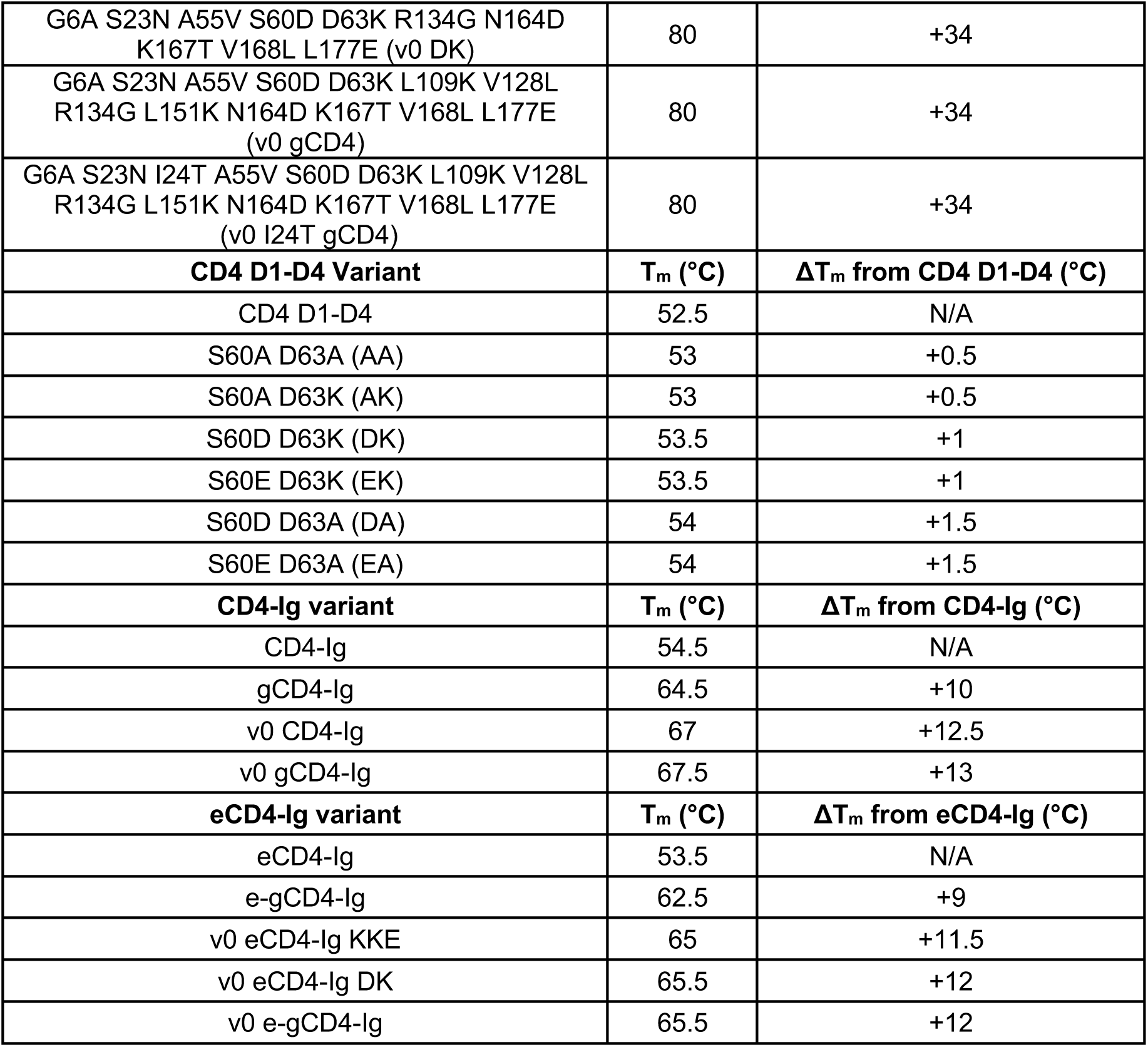
Apparent melting temperatures of CD4-containing constructs. Samples were measured in technical triplicate and the averages are reported.

**Extended Data Table 2:**
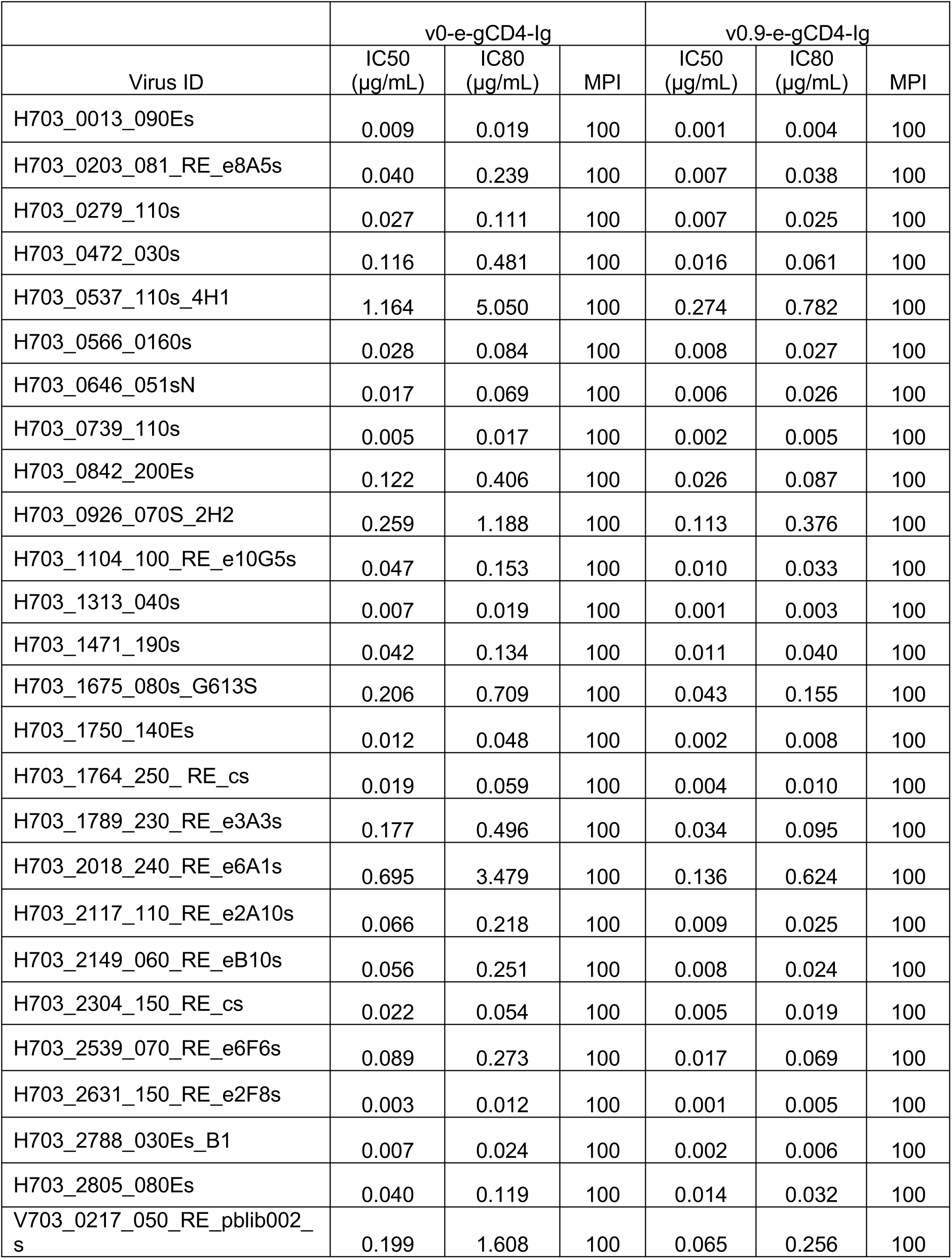
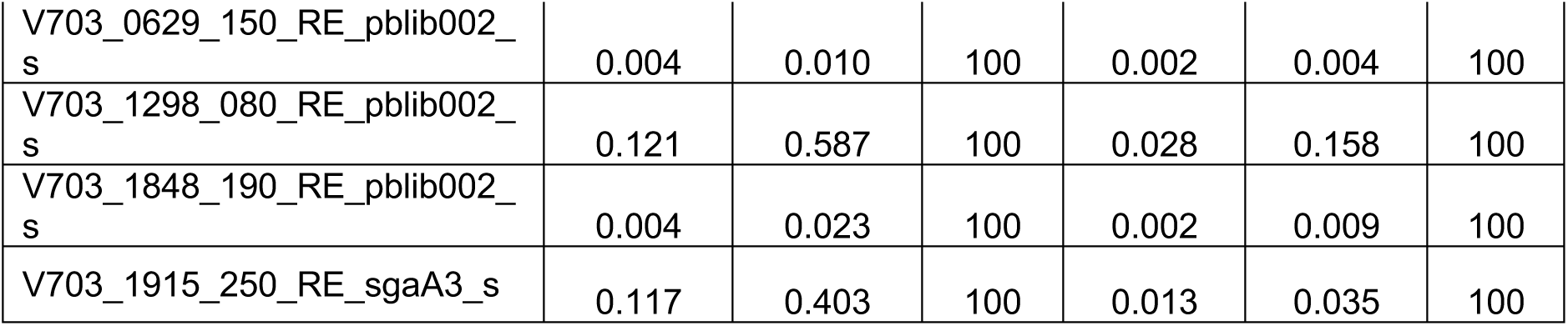
IC_50_, IC_80_, and MPI (maximum percent inhibition) values of v0-egCD4-Ig and v0.9-e-gCD4-Ig against 30 AMP trial pseudoviruses.

**Extended Data Fig. 1:**
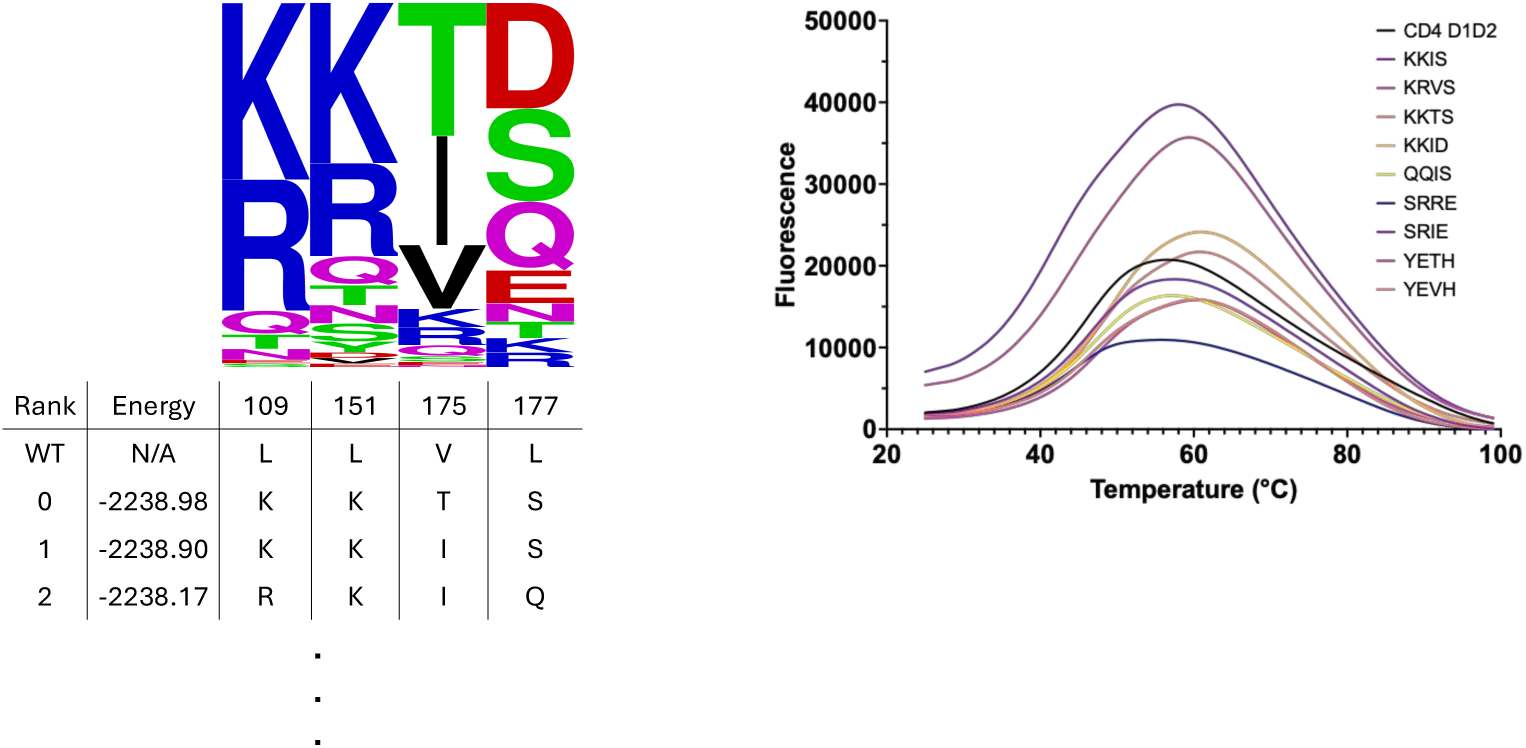
Representative energy calculation of the D2 domain hydrophobics and DSF data. Shown is an example of an output from TRIAD using the PHOENIX scoring function and a sequence logo of the suggested amino acids at each position. The DSF data of nine variants is shown and their T_m_s are reported in **Extended Data Table 1**.

**Extended Data Fig. 2:**
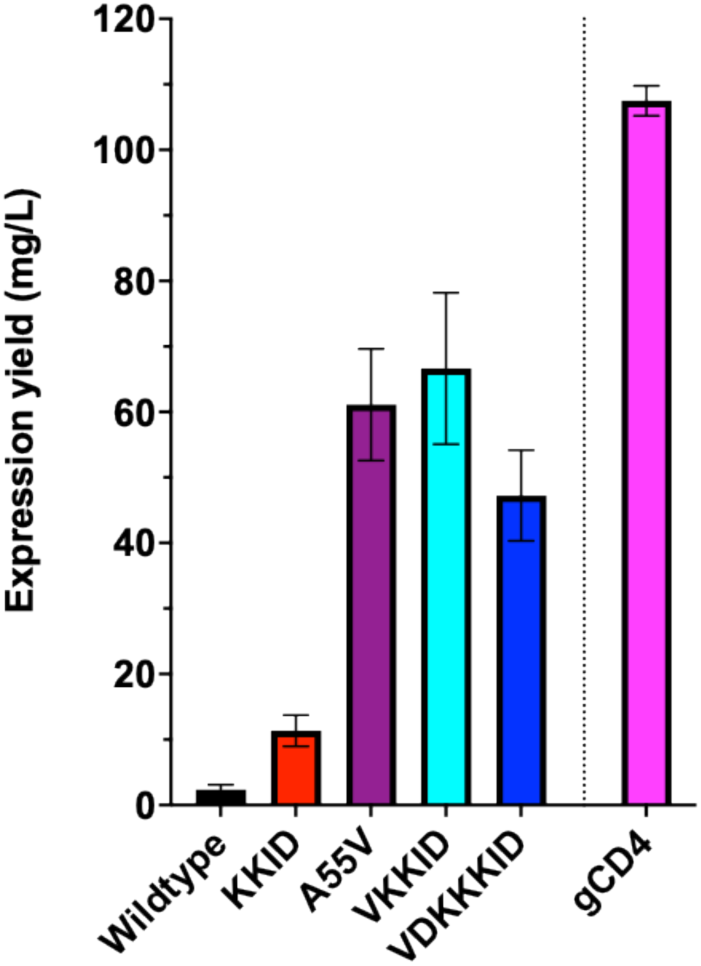
Expression yield of CD4 D1D2 variants in Expi293 cells. Proteins were expressed in 20 mL volumes and purified as described in the Methods in technical triplicates. Yields were then calculated for one liter expressions assuming a linear relationship with culture volume. gCD4 contains the VDKKKE substitutions and was expressed in the p3BNC plasmid. All other proteins were expressed in pTT5. Error bars are s.d.

**Extended Data Fig. 3:**
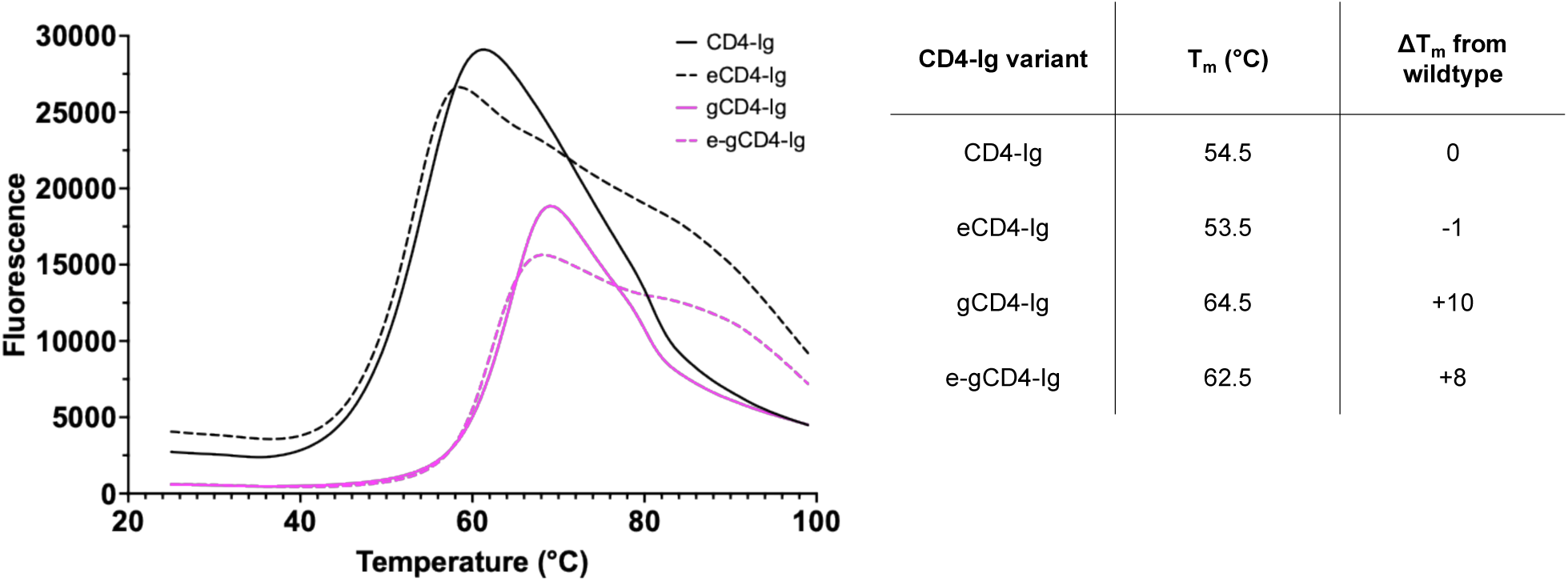
Apparent melting temperatures of CD4-Ig variants. Curves represent the average of technical triplicates.

**Extended Data Fig. 4:**
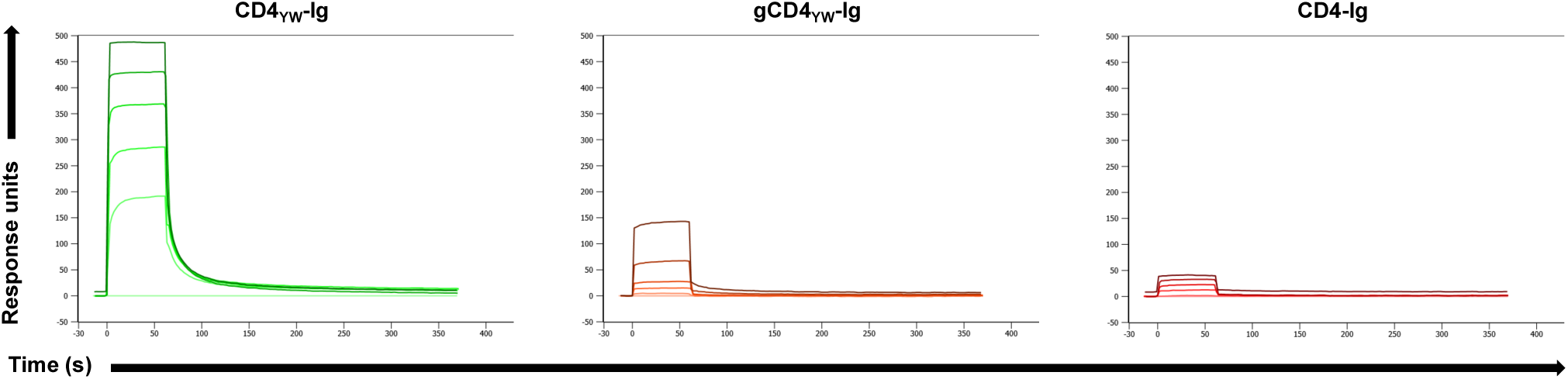
SPR of CD4-Ig variants against immobilized HLA-DR1. Proteins were injected for multi-cycle kinetics using 3-fold dilutions series from 50 μM to 0.62 μM as in **Fig. 2**.

**Extended Data Fig. 5:**
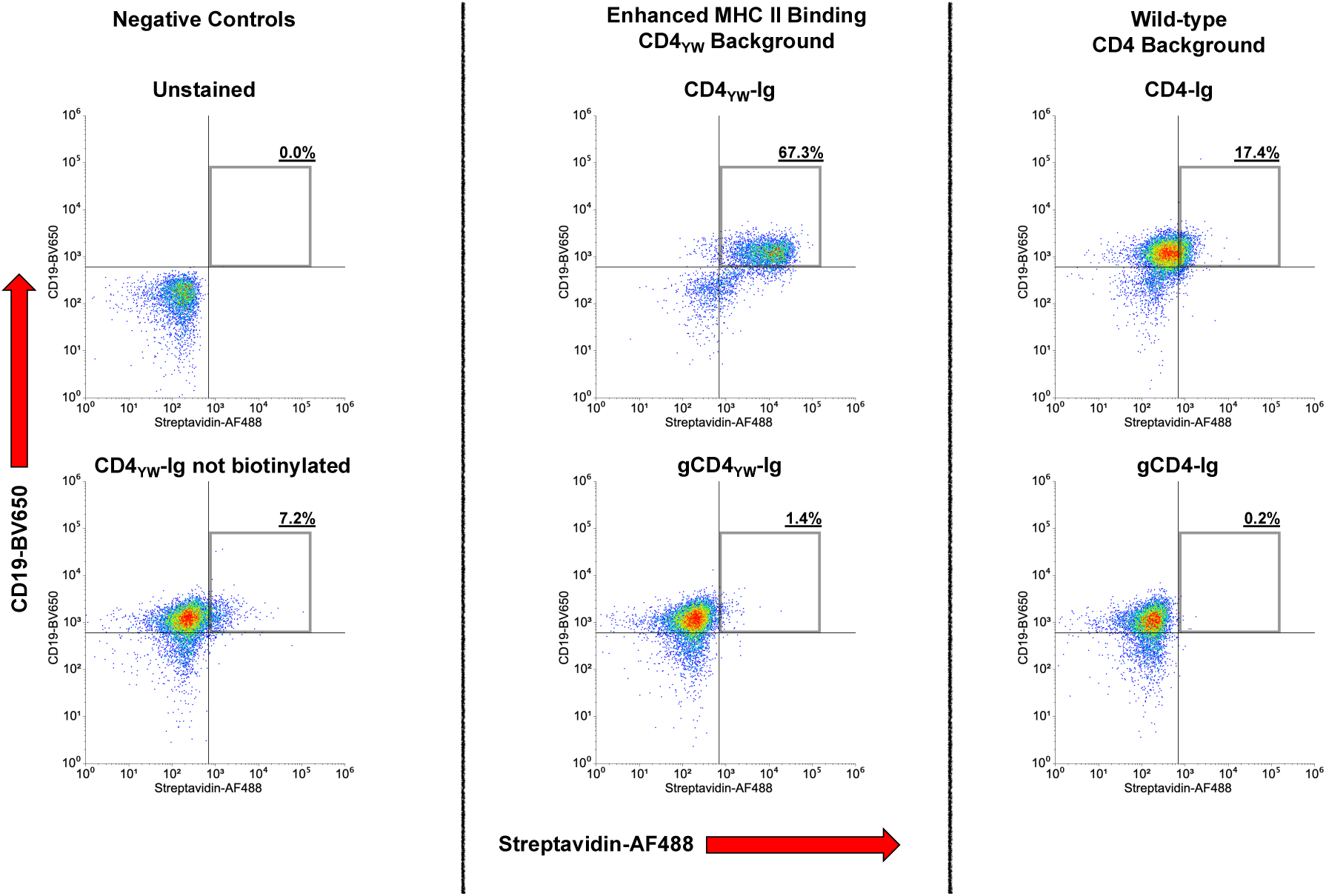
Flow cytometry of CD4-Ig variants using tonsillar cells from donor #667. Flow cytometry was performed as in **Fig. 3**.

**Extended Data Fig. 6:**
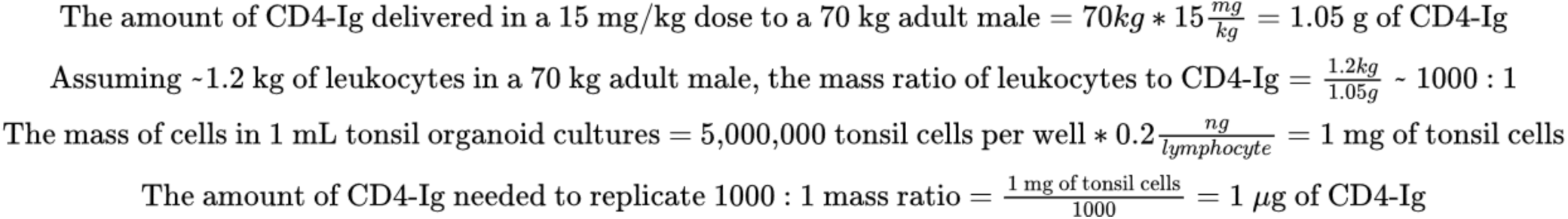
Calculation of the CD4-Ig amount needed to mimic clinical trial dosing in tonsil organoid cultures. Leukocyte mass was assumed using a reference 70 kg adult male^52^.

**Extended Data Fig. 7:**
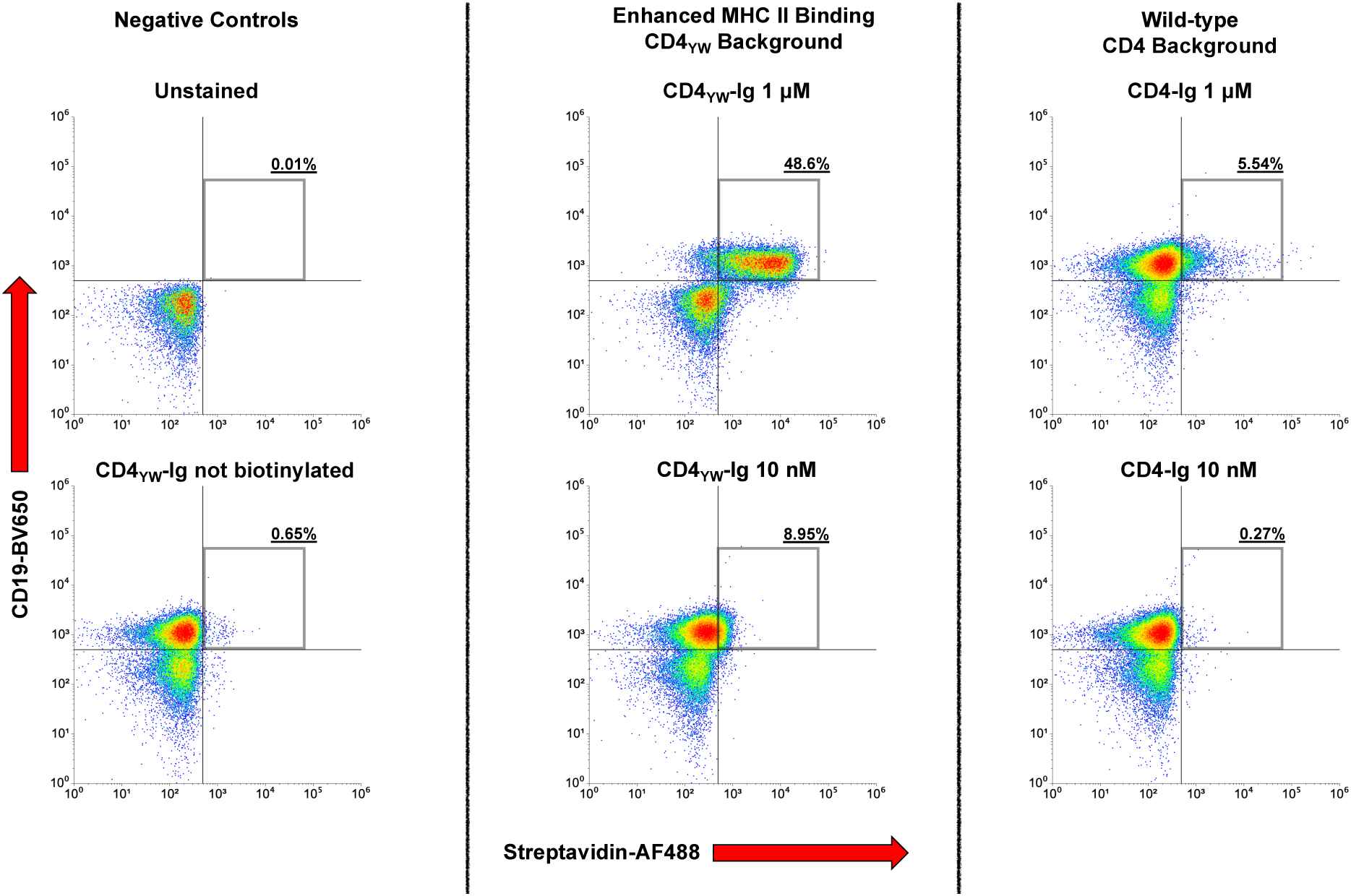
Flow cytometry of CD4-Ig variants at differing concentrations. In order to mimic clinical trial dosing in tonsil organoid cultures, CD4-Ig needed to be used at 10 nM (as determined in **Extended Data Fig. 6**). CD4_YW_-Ig and CD4-Ig at 1 μM and 10 nM were tested to determine which construct should be used in tonsil organoid cultures at 10 nM. Tonsil cells are from donor #667.

**Extended Data Fig. 8:**
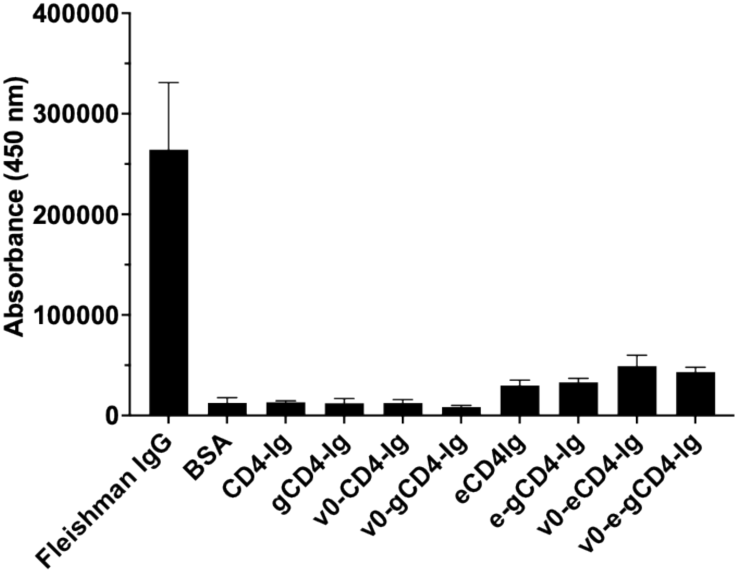
Baculovirus polyreactivity assay of CD4-Ig variants. Proteins were assayed in quadruplicate. Fleischmann IgG was used as a positive control and BSA as a negative control. Error bars are s.d.

**Extended Data Fig. 9:**
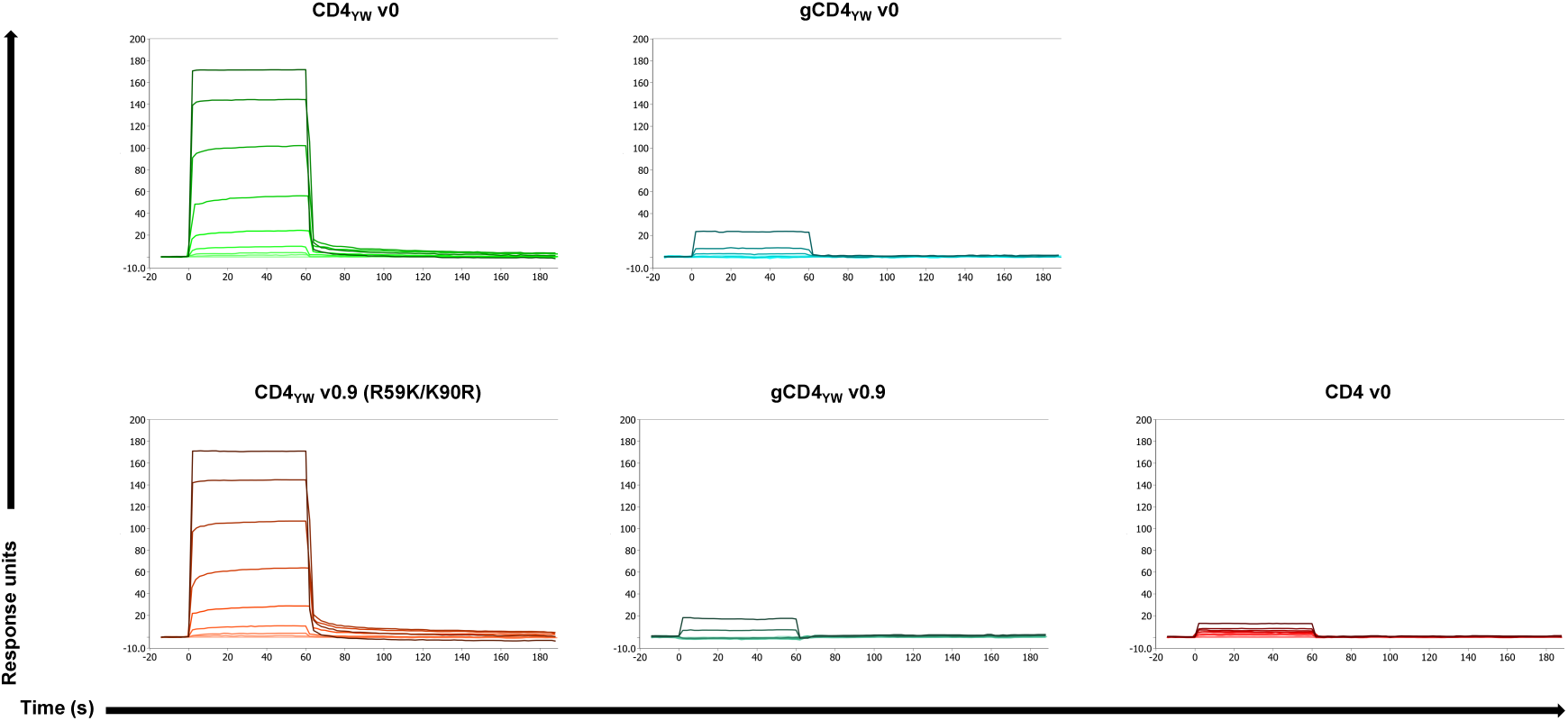
SPR of new CD4 variants against HLA-DR1. SPR was performed using multi-cycle kinetics and a 3-fold dilution series from 50 μM to 0.023 μM as in **Fig. 2**.

